# Excess ribosomal protein production unbalances translation in Fragile X Syndrome

**DOI:** 10.1101/2022.04.07.487442

**Authors:** Sang S. Seo, Susana R. Louros, Natasha Anstey, Miguel A. Gonzalez-Lozano, Callista B. Harper, Nicholas C. Verity, Owen Dando, Sophie R. Thomson, Jennifer C. Darnell, Peter C. Kind, Ka Wan Li, Emily K. Osterweil

**Author notes:** **Lead Contact:** Emily K. Osterweil, University of Edinburgh, Hugh Robson Building, George Square, Edinburgh, EH8 9XD, UK. Equal contributions (Sang S. Seo, Susana R. Louros).

## Abstract

Dysregulated protein synthesis is a core pathogenic mechanism in Fragile X Syndrome (FX). The mGluR Theory of FX predicts that pathological synaptic changes arise from the excessive translation of mRNAs downstream of mGlu_1/5_ activation. Here, we use a combination of CA1 pyramidal neuron-specific TRAP-seq and proteomics to identify the overtranslating mRNAs supporting exaggerated mGlu_1/5_-induced long-term synaptic depression (mGluR-LTD) in the FX mouse model (*Fmr1^-/y^*). Surprisingly, our results identify a robust translation of ribosomal proteins (RPs) upon mGlu_1/5_ stimulation that coincides with a reduced translation of long mRNAs encoding synaptic proteins. These changes are mimicked and occluded in *Fmr1^-/y^* neurons. Inhibiting RP translation significantly impairs mGluR-LTD and prevents the length-dependent shift in the translating population. Together, these results suggest that pathological changes in FX result from a length-dependent alteration in the translating population that is supported by excessive RP translation.

## Introduction

Several genes involved in mRNA translation have been identified as penetrant risk factors for autism ^1,2^. Most prevalent is the *FMR1* gene encoding the RNA binding protein Fragile X Mental Retardation Protein (FMRP)^3^. In FX, excessive neuronal protein synthesis is a core phenotype that is conserved from fly to humans ^4–7^, and it is believed to be pathological because targeting translation control signaling pathways can correct electrophysiological and behavioral deficits in *Fmr1^-/y^* animal models ^4–6,8–10^. FMRP is an RNA binding protein that has been implicated in the repression of translation. A leading mechanistic model proposes that FMRP slows elongation of target transcripts, resulting in a depression of the translation of these targets in FX neurons ^3^. Recent studies using ribosome profiling and RNA-seq have provided important support for this model, showing that there is a reduction in ribosome pausing along long mRNAs in *Fmr1^-/y^* brain ^11,12^. However, the way in which this dysregulation of protein synthesis results in neurological disruptions in FX has not been established. This is due in large part to the lack of information about the mRNAs over-translated in *Fmr1^-/y^* brain that are impeding synaptic function.

A key synaptic phenotype in the *Fmr1^-/y^* model is an exaggeration of long-term synaptic depression induced by activation of group I metabotropic glutamate receptors (mGluR-LTD) in hippocampal CA1, a major form of synaptic plasticity that contributes to learning ^13,14^. This phenotype has been observed in numerous studies of *Fmr1^-/y^* mice and rats, and has been used as a model for synaptic dysfunction in FX ^7^,^15^. Inhibition of protein synthesis specifically in CA1 pyramidal (pyr) neurons blocks mGluR-LTD, showing that de novo translation in these neurons is required for plasticity ^16^. In the *Fmr1^-/y^* hippocampus, LTD is not only greater in magnitude, it is also persistent in the presence of protein synthesis inhibitors that disrupt mGluR-LTD in WT ^17^. Based on these results and others, The mGluR Theory of FX was proposed as a conceptual framework for interpreting how the loss of FMRP-regulated protein synthesis leads to pathological changes in the FX brain ^14^. According to this model, protein synthesis downstream of group 1 mGluRs (mGlu_1/5_) is unchecked in the absence of FMRP, causing an exaggeration of mGluR-LTD and other mGlu_1/5_-associated changes. Multiple studies have shown that inhibition of mGlu_5_, the predominant group 1 mGluR in forebrain, corrects pathological changes in *Fmr1^-/y^* models, including excessive protein synthesis, altered dendritic spine function, exaggerated LTD, epileptiform activity, learning deficits, and expression of audiogenic seizures ^8,18^. Identifying the proteins that are both over-synthesized in *Fmr1^-/y^* and synthesized in response to mGlu_5_ activation thus represents a unique opportunity to assess how altered translation in FX neurons disrupts synaptic function. Although there have been important candidate-driven analyses of this question ^19,20^, there has yet to be a comprehensive cell-type specific analysis of the mRNAs translated in response to LTD, and analysis of the way that these are altered in the *Fmr1^-/y^* hippocampus.

In this study, we identified mRNAs both over-translated in *Fmr1^-/y^* and translated in response to mGluR-LTD stimulation in CA1 pyr neurons using a combination of cell type-specific Translating Ribosome Affinity Purification and RNA-seq (TRAP-seq) and label-free proteomics. Surprisingly, we find that the most commonly over-translated and over-expressed population in *Fmr1^-/y^* neurons is comprised of ribosomal proteins (RPs). Using multiple methodologies, we confirm a significant per-cell increase in the expression of RPs and ribosomal RNA (rRNA) in *Fmr1^-/y^* neurons, which has not been previously described. Furthermore, this first translation profiling analysis of mGluR-LTD in FX reveals a similar significant increase in RP translation that is occluded in *Fmr1^-/y^* neurons. In fact, the majority of translation changes induced by mGluR-LTD in WT are strikingly similar to those constitutively changed in *Fmr1^-/y^* neurons, providing strong support for the mGluR Theory. Interestingly, further analysis shows that the RP translation that is induced by mGluR-LTD and is constitutive in *Fmr1^-/y^* neurons alters the translating population in a manner that reduces translation of longer-length transcripts. This results in a reduction in the expression of proteins involved in synaptic stability encoded by these long transcripts, including FMRP targets. Incubation with the ribogenesis inhibitor CX-5461 prevents the reduced translation of long mRNAs by mGlu_1/5_ stimulation and impairs mGluR-LTD, directly linking the translation changes observed in CA1 neurons to a hallmark FX phenotype. Together, these results show that the composition of the translating mRNA population in *Fmr1^-/y^* neurons is comprised of an overabundance of RPs and a paradoxical deficit in long mRNAs encoding synaptic proteins, which mimics the translation downstream of mGlu_1/5_ activation. This would suggest a reconceptualization of the relationship between excess translation and pathological synaptic changes in FX.

### Ribosomal proteins are over-translated in *Fmr1^-/y^* neurons

Identification of the over-translating mRNAs in FX is essential for understanding how synaptic pathology emerges. In previous work, we investigated ribosome-bound mRNAs in *Fmr1^-/y^* hippocampal CA1 pyr neurons using TRAP-seq, which revealed over-translated mRNAs that compensate for the loss of FMRP ^21^. While this study identified a novel target that can be positively modulated to correct *Fmr1^-/y^* phenotypes, it did not reveal an obvious group of targets driving the pathology of FX. To identify this group, we tried a new approach comparing the mistranslating mRNAs identified by TRAP to the synapse-enriched proteome of the *Fmr1^-/y^* hippocampus (**Fig. 1a**). We reasoned that a comparison of the population both over-translated and over-expressed, in the same hippocampal slice preparation, would more accurately identify contributors to pathological changes.

**Figure 1.**
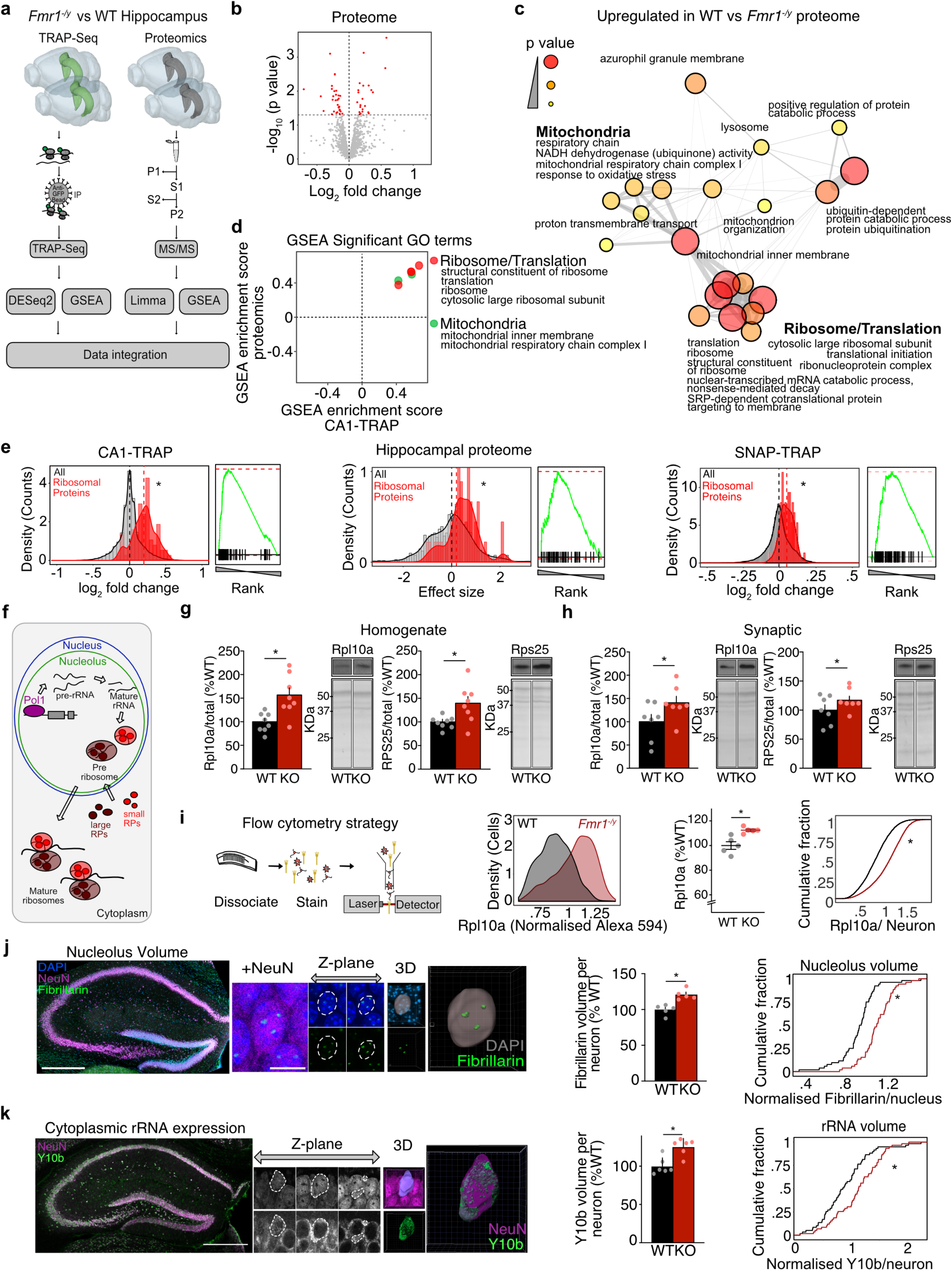
Ribosomes are over-produced in *Fmr^-/y^* hippocampal neurons. **(a)** Schematic of the experimental strategy. Multi-Omics approach was used to identify over-translating and over-expressed proteins. **(b)** Volcano plot of proteomics analysis of hippocampal P2 fractions isolated from WT/*Fmr1^-/y^* littermates. Significant targets (p < 0.05) are denoted in red. **(c)** GSEA analysis of the proteomics dataset identified overexpression of ribosome/translation and mitochondrial related GO terms in the *Fmr1^-/y^* (adjusted p value < 0.1). **(d)** Overlap of significantly upregulated gene sets identified by GSEA in both TRAP-seq and proteomic datasets reveals ribosome/translation and mitochondrial GO terms as the most enriched in the upregulated populations. **(e)** Analysis of the 80 RP population versus total population in *Fmr1^-/y^* versus WT CA1-TRAP, proteomic, and SNAP-TRAP datasets reveal a significant upregulation in all 3 datasets (two sample z test: z = 6.54, *p = 6.25X10^-11^, z = 2.51, *p = 0.012, z = 5.12, *p = 3.11X10^-7^, CA1-TRAP, P2 proteomic, Snap-TRAP respectively). **(f)** Schematic of ribogenesis showing precursor rRNA transcription in the nucleolus by RNA Pol I, followed by splicing and folding into mature rRNA subunits, RP production, and association with mature ribosomes upon export to the cytoplasm. **(g)** Immunoblotting of hippocampal homogenates from *Fmr1^-/y^* and WT littermates revealed a significant increase in the large ribosome-associated protein Rpl10a (WT = 100 ± 7.110%, *Fmr1^-/y^*= 158.7 ± 15.19%, *p = 0.0008, N = 8) and the small ribosome-associated protein Rps25 (WT = 100 ± 4.528%, *Fmr1^-/y^*= 139.7 ± 14.28%, *p = 0.0468, N= 8). **(h)** Synaptoneurosomes isolated from WT and *Fmr1^-/y^* hippocampi show the same increase in Rpl10a (WT = 100 ± 15.44%, *Fmr1 ^-/y^*= 140.4 ± 13.95%, *p = 0.0436, N= 7) and Rps25 (WT = 100 ± 9.178%, *Fmr1^-/y^*= 117.1 ± 7.098%, *p = 0.0235, N= 7) **(i)** Schematic showing steps for FACS immunostaining. Comparison of CA1 neurons isolated from 5 littermate pairs shows a significant increase in Rpl10a expression in *Fmr1^-/y^* vs WT (paired t test, WT = 100 ± 3.1 %, *Fmr1^-/y^* = 112.5 ± 1.08 %, *p = 0.02915, KS test *p < 0.0001) **(j)** Sections from *Fmr1^-/y^* and WT littermate brains were immunostained for NeuN, fibrillarin and DAPI or NeuN and Y10b before confocal imaging of the dorsal hippocampal CA1 region (scale = 10 μm). Total nucleolar volume was quantified from 3D reconstruction of fibrillarin staining. Analysis shows a significant increase in total nucleolar volume per cell in *Fmr1^-/y^* vs WT neurons when quantified per animal (paired t test, WT = 100 ± 3.277 %, *Fmr1^-/y^*= 121.2 ± 3.277 % * p = 0.032 N= 5 littermate pairs) or as a cumulative distribution of all neurons (KS test *p < 0.0001, N = 52 neurons). **(k)** Volume of rRNA was quantified from 3D reconstruction of Y10b staining normalized to NeuN. A significant increase in rRNA volume was observed in in *Fmr1^-/y^* neurons (paired t test, WT = 100 ± 4.458 %, *Fmr1^-/y^*= 125.4 ± 4.458 %, * p = 0.0358, N= 6 littermate pairs, KS test *p = 0.00792, N = 59 neurons).

To identify proteomic changes, we isolated hippocampal slices from 5 pairs of littermate WT and *Fmr1^-/y^* mice of a juvenile age (P25-32), which is consistent with the CA1 TRAP-seq samples isolated from 6 pairs of juvenile *Fmr1^-/y^* and WT hippocampus. Synapse-enriched P2 fractions were isolated from each set of slices. This latter step is regularly used when examining proteomic changes relevant to synaptic function, and it is necessary to isolate the synapse-relevant proteome from the bulk tissue that contains cell bodies from all cell types including glia ^22,23^. Isolated fractions were processed for label-free Mass Spectroscopy (MS) and differential expression was determined as described in Methods. Consistent with previous proteomic studies of *Fmr1^-/y^* cortex, we find that the changes in the steady-state proteome of the *Fmr1^-/y^* hippocampus are modest, with 52 proteins identified as significantly altered (**Fig. 1b, Table 1**). We therefore further interrogated this dataset using a rank-based Gene Set Enrichment Analysis (GSEA), which was developed to identify gene sets that shift together ^24^. This analysis revealed that 20 gene sets were significantly upregulated in the *Fmr1^-/y^* proteome (**Table 2**). Surprisingly, the most significantly upregulated gene sets were related to ribosomes (**Fig. 1c**). Next, we performed the same GSEA on the CA1-TRAP dataset collected in our previous study, identifying 14 gene sets significantly upregulated ^21^ (**Supplementary Fig. 1a**, **Table 2**). Remarkably, a comparison of gene sets significantly upregulated in both proteomic and CA1-TRAP populations again identified ribosomes (**Fig. 1d**).

To determine whether the GSEA results reflected an actual increase in ribosomes, we began by assessing the 80 distinct ribosomal proteins (RPs) associated with eukaryotic ribosomes ^25^. Although recent work has discovered that ribosomes can exhibit a specialized composition of RPs, the majority of ribosomes are believed to express RPs in a stoichiometric fashion ^26^. We therefore analyzed the expression of the 80 RP gene set as a group. These results show a significant upregulation of RPs in both the CA1 TRAP and proteome of *Fmr1^-/y^* hippocampus (**Fig. 1e**). Quantifying the ribosome-bound mRNAs isolated by TRAP can reflect a difference in transcript abundance as well as translation. To determine whether a change in the abundance of RP transcripts could account for the change seen in the TRAP fraction, we normalized the counts obtained from the TRAP samples to those from the total hippocampal transcriptome. Our results show that although there is a significant increase in RP expression in the total *Fmr1^-/y^* mRNA population, the increase in the TRAP-bound fraction is greater and persists after normalization to this total (**Supplementary Fig. 1b**).

Although our investigation was largely focused on CA1 pyr neurons due to the pivotal role these cells play in mGluR-LTD, we wondered whether the increase in RP expression could be seen in the broader neuronal population. To determine this, we performed TRAP-seq on hippocampi isolated from 4 littermate WT and *Fmr1^-/y^* pairs expressing EGFP-L10a under the *Snap25* promoter, which exhibits pan-neuronal expression throughout postnatal development (**Supplementary Fig. 1c, Table 3**). Similar to our results from CA1, we find RP transcripts are upregulated in the *Fmr1^-/y^* SNAP-TRAP fraction (**Fig. 1e, Supplementary Fig. 1d**). This shows increased RP translation is expressed in the broader *Fmr1^-/y^* neuron population.

### Ribosomes are over-produced in *Fmr1^-/y^* neurons

Although surprising, a confirmed increase in ribosome production in *Fmr1^-/y^* neurons would have important implications for understanding the protein synthesis dysregulation seen in FX. Ribogenesis is a complex process that involves both transcription of rRNA subunits in the nucleolus, and production of RPs that ultimately assemble into mature ribosomes that are exported into the cytoplasm (**Fig. 1f**). To determine whether the observed increase in RP translation reflected an increase in ribogenesis in *Fmr1^-/y^* neurons, we performed multiple validation experiments. First, we performed immunoblotting for RPs associated with both small and large ribosomal subunits and found both the large subunit-associated Rpl10a and small subunit-associated Rps25 are significantly overexpressed in *Fmr1^-/y^* versus WT hippocampus (**Fig. 1g, Supplementary Fig. 1e**). The same increase is also seen in synaptoneurosome fractions (**Fig. 1h**). Next, it was essential to determine whether the increased RP expression could be seen on a per-neuron basis to rule out the possibility that our results are due to a difference in cell size or composition. To assess per-neuron expression, we immunostained dissociated neurons from *Fmr1^-/y^* and WT hippocampi for Rpl10a and quantified per-cell expression using flow cytometry (**Fig. 1i**). Analysis of 6 littermate pairs reveal a significant increase in the expression of Rpl10a in individual *Fmr1^-/y^* neurons. These results provide strong validation of the results seen in our CA1-TRAP and proteomic datasets.

Along with RPs, functional ribosomes require the presence of 4 core rRNA subunits, which are transcribed in the nucleolus, which increases in size during periods of high ribogenesis ^27^. To verify that expression of the rRNA core was elevated in *Fmr1^-/y^* versus WT neurons, we first measured nucleolar volume by immunostaining for the nucleolar marker fibrillarin (**Fig. 1j**). Analysis of reconstructed confocal z-stacks reveals that nucleolar volume is significantly increased in *Fmr1^-/y^* neurons measured as either an increase in total nucleolar volume per neuron (**Fig. 1j**) or as the average volume of nucleoli per neuron (**Supplementary Fig. 1f)**. To confirm that the amount of mature rRNA outside the nucleolus was also increased, we performed immunostaining using a Y10b antibody raised against rRNA ^28^. Calculation of Y10b staining per volume of NeuN shows a significant increase in rRNA expression in *Fmr1^-/y^* versus WT hippocampal neurons (**Fig. 1k**), with no significant difference in the total volume of NeuN (**Supplementary Fig. 1f**). Together, our results confirm an elevated expression of ribosomes in *Fmr1^-/y^* neurons.

### mGluR-LTD induces increased RP translation and decreased translation of synaptic proteins, which is mimicked and occluded in *Fmr1^-/y^*

The mGluR Theory suggests that the mRNAs translated in response to mGlu_5_ activation is already over-translating in *Fmr1^-/y^* neurons, contributing to the protein synthesis-independence of LTD (**Fig. 2a**). Supporting this, protein synthesis downstream of mGlu_1/5_ activation is saturated in *Fmr1^-/y^* CA1 ^4,29^. Although previous work has examined proteomic changes induced by the mGlu_1/5_ agonist DHPG in cultured neurons ^30^ and hippocampal slices ^31^, a translation profiling analysis of neurons expressing LTD has not been performed. To assess this, we performed TRAP on hippocampal slices prepared from WT and *Fmr1^-/y^* littermate CA1-TRAP mice stimulated with DHPG using a protocol that induces mGluR-LTD (**Fig. 2b**). TRAP-seq was performed at 30 min post-stimulation, which is consistent with the timeframe of protein-synthesis dependent LTD ^32^. Our results show that mGluR-LTD has a striking impact on the translating population in WT CA1 neurons, causing the differential expression of 696 transcripts in the TRAP population (**Fig. 2c, Supplementary Fig. 2a, Table 4**). In contrast, only 15 transcripts are significantly differentially expressed in the *Fmr1^-/y^* CA1-TRAP after DHPG stimulation. These results are consistent with evidence showing that DHPG does not stimulate translation in *Fmr1^-/y^* hippocampus, presumably due to a saturation of the translation machinery ^4,29^. This effect in *Fmr1^-/y^* neurons is not due to a general disruption in signalling downstream of mGlu_1/5_ as the activity-driven upregulation in the immediate early genes *Npas4* and *Arc* is seen in both genotypes (**Fig. 2d**), an effect that is validated by qPCR analysis of additional DHPG-treated slices (**Supplementary Fig. 2b**) ^33,34^.

**Figure 2.**
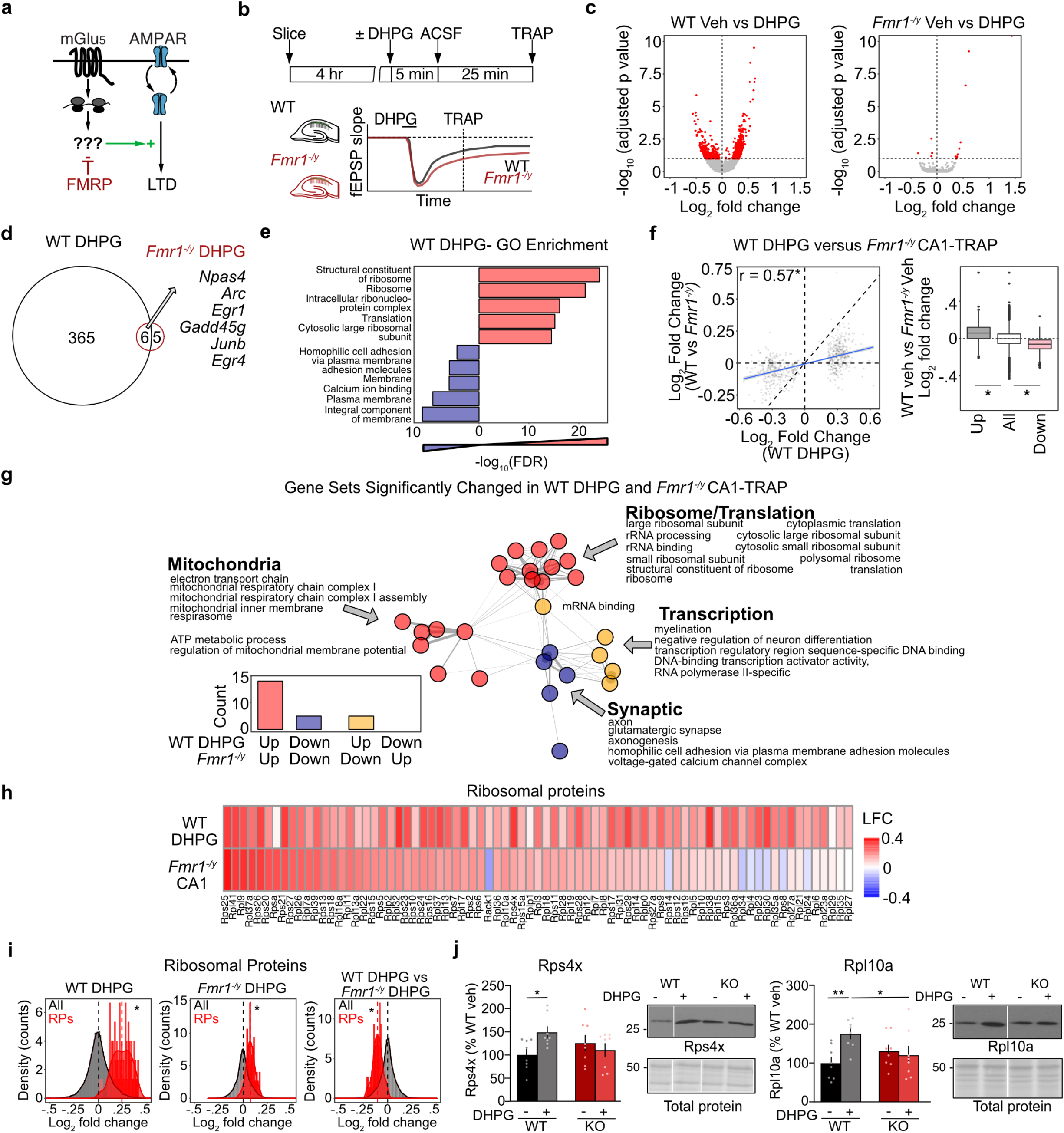
mGluR-LTD induces translation changes that are mimicked and occluded in *Fmr1^-/y^* neurons, including translation of RPs. **(a)** Schematic of the mGluR Theory of FX. (**b**) Schematic of the experimental strategy. WT and *Fmr1^-/y^* slices were recovered and stimulated for mGluR-LTD using a protocol of 5 min DHPG followed by a washout of 25 mins, after which TRAP was performed. **(c)** Volcano plots of TRAP-seq data show that DHPG induces substantial significant translational changes in WT but not in *Fmr1^-/y^* CA1 neurons (DESeq2 adjusted p-value < 0.1). **(d)** Quantification shows 371 targets upregulated in WT and only 11 targets in *Fmr1^-/y^*. Overlapping include immediate early genes that report neuronal activity, including *Npas4* and *Arc*. **(e)** DAVID GO enrichment analysis of the up- and downregulated populations induced by DHPG reveals that ribosome/translation related transcripts are enriched in the upregulated population whereas membrane/ calcium ion binding transcripts are enriched in the downregulated fraction. **(f)** Transcripts significantly changed in WT DHPG are significantly correlated with basal expression changes in *Fmr1^-/y^* (r = 0.57, *p < 2.2X10^-16^). Analysis of the significantly up- and downregulated transcripts in the WT DHPG dataset shows they exhibit a significant basal log_2_ fold change difference in the *Fmr1^-/y^* population as well when compared to the total population (Kruskal-Wallis test *p < 2.2 X10^-16^, post-hoc Wilcoxon rank sum test up *p < 2.2X10^-16^, down *p <2.2X10^-16^). **(g)** To determine whether the gene sets altered with LTD are similar to those already altered in the *Fmr1^-/y^* translating population, GSEA was performed on the WT DHPG population and significantly changed gene sets (adjusted p-value < 0.1) were compared to those significantly changed in the *Fmr1^-/y^* population (p-value < 0.01). This reveals a striking overlap with ribosome/mitochondrial terms upregulated in both WT DHPG and *Fmr1^-/y^*, and synaptic terms downregulated in both populations. (**h**) A heatmap of log2 fold change shows that RPs are basally upregulated in *Fmr1^-/y^* and in WT after DHPG stimulation. **(i)** RPs show an increase with DHPG in WT CA1-TRAP that is seen to a lesser degree in *Fmr1^-/y^* CA1-TRAP (z test WT-LTD: z = 14.74, *p < 2.2X10^-16^, *Fmr1^-/y^-LTD*: z = 6.35, *p = 2.2X10^-10^). Comparison of the DHPG effect on RP expression shows the response in *Fmr1^-/y^* is occluded when compared to WT (z test: z =-8.68, *p < 2.2X10^-16^). **(j)** Immunoblot analysis from synaptoneurosome fractions isolated from WT and *Fmr1^-/y^* slices stimulated with DHPG shows a significant upregulation of Rps4x (WT = 100 ± 11.6%, WT DHPG = 148.9 ± 11.9%, *Fmr1^-/y^* = 125.6 ± 16.4%, *Fmr1^-/y^* DHPG = 110.4 ± 14.0%. 2-ANOVA genotype x treatment *p = 0.0258, WT vs WT DHPG p= 0.0169. N= 8) and Rpl10a (WT = 100 ± 14.8%, WT DHPG = 175.3 ± 12.9%, *Fmr1^-/y^* = 130.9 ± 15.1%, *Fmr1^-/y^* DHPG = 120.6 ± 22.8%. 2-ANOVA genotype x treatment *p = 0.0212, WT vs WT DHPG p=0.0149. N= 8) in WT slices and no change in *Fmr1^-/y^*.

To further explore these changes, we performed GO analyses on the significantly up- and downregulated transcripts stimulated with DHPG in the WT CA1-TRAP. This revealed a striking functional divergence in the populations increased and decreased in translation (**Fig. 2e**). Interestingly, the mRNAs most enriched in the WT DHPG upregulated population were related to ribogenesis, similar to what is seen in the *Fmr1^-/y^* CA1-TRAP. In contrast, the population significantly reduced in translation with DHPG is enriched for transcript encoding ion channels, cell adhesion molecules and structural synaptic elements. GSEA reveals the same functional divergence is seen in the up- and downregulated populations (**Supplementary Fig. 2c-d**).

Next, to determine whether the translation changes induced by mGluR-LTD in WT are similar to the basal translation changes in the *Fmr1^-/y^* CA1, we compared the expression of the LTD transcripts in DHPG-treated WT slices to the CA1-TRAP population isolated from *Fmr1^-/y^* hippocampus. Analysis of the transcripts significantly changed with DHPG in WT shows a clear significant correlation to what is basally altered in the *Fmr1^-/y^* CA1-TRAP (**Fig. 2f**). In addition, analysing the expression of significantly up- and down-regulated populations in the WT DHPG population shows that these are similarly up- and downregulated in the unstimulated *Fmr1^-/y^* population as compared to the average. To explore the functional relevance of the translation changes induced by DHPG and occluded in *Fmr1^-/y^* neurons, we performed GSEA to compare gene sets significantly altered with DHPG in WT versus those basally altered in the *Fmr1^-/y^* CA1-TRAP. This revealed a striking overlap, with 82% of gene sets significantly changed with DHPG in WT also significantly changed in the same direction in the basal *Fmr1^-/y^* population (**Fig. 2g, Table 5**). Interestingly, the population that is both upregulated with DHPG and constitutively upregulated in *Fmr1^-/y^* is enriched for RPs and rRNA processing, as well as mitochondrial function. In contrast, the downregulated population is enriched for glutamatergic synapse, homophilic cell adhesion, voltage-dependent calcium channels and axonal function. These results indicate that changes in the translating fraction of CA1 neurons that occur during mGluR-LTD in WT are largely mimicked and occluded in the *Fmr1^-/y^* hippocampus, supporting the model put forth by the mGluR Theory.

To identify the newly-synthesized proteins most likely involved in the exaggerated LTD phenotype, we examined the targets significantly upregulated in DHPG-treated WT slices and those basally upregulated in the *Fmr1^-/y^* CA1-TRAP population at a threshold of p < 0.05. This analysis shows a significant overlap between the upregulated populations and identified 92 candidates (**Supplementary Fig. 2e, Table 6**). Consistent with our GSEA results, ribosome-related transcripts comprise the greatest proportion of the overlapping population (**Supplementary Fig. 2f**). Analysis of individual RPs reveals that most are upregulated in both datasets, with 11 of the 12 RPs significantly upregulated in the *Fmr1^-/y^* CA1-TRAP population also significantly upregulated with DHPG in WT (**Fig. 2h, Table 7**). This surprising result led us to question whether new RP synthesis was supporting mGluR-LTD. Although canonical ribogenesis requires transcription of rRNA in the nucleus, several studies have shown that RP transcripts are abundant in axons and dendrites, and that RPs are locally synthesized ^35–38^. The functional relevance of new RP synthesis in neurons is shown in Shigeoka et al., who provide strong evidence that local synthesis of Rps4x is critical for axon branching ^39^. More recently, Fusco et al. show that RP transcripts in dendrites are rapidly synthesized and incorporated into existing ribosomes^35^. Both studies show that a subset of RPs proximal to the cytosolic face of the ribosome are found to be more frequently incorporated into existing ribosomes. Analysis of our CA1-TRAP results shows that 6 of the 11 rapidly-exchanging RPs identified by Fusco et al. and 9 of the 11 identified by Shigeoka et al. are significantly increased with DHPG (**Table 7**). Interestingly, Rps4x is one of the most significantly increased RPs in the DHPG dataset.

To determine whether the upregulation of RPs with DHPG was occluded in *Fmr1^-/y^* CA1 neurons, we quantified the RP gene set in both genotypes after stimulation. Our results show that DHPG significantly increases the RP gene set in both WT and *Fmr1^-/y^* neurons, however the effect in the *Fmr1^-/y^* population is significantly impaired when compared to that seen in WT (**Fig. 2i**). To determine whether this effect could be seen at the protein level, we isolated synaptoneurosome fractions from WT and *Fmr1^-/y^* slices 1 hour after stimulation with DHPG and performed immunoblotting for individual RPs. Our results show that DHPG simulation significantly increases both Rpl10a and Rps4x in the synaptic fractions of WT slices, and these RPs are basally elevated in *Fmr1^-/y^* slices (**Fig. 2j, Supplementary Fig. 2g)**. These results confirm that there is a rapid increase in RPs at WT synapses after DHPG treatment that is mimicked and occluded in the *Fmr1^-/y^* hippocampus.

### Inhibiting RP translation blocks mGluR-LTD in WT but not *Fmr1^-/y^* hippocampal CA1

Our TRAP-seq results showing the significant increase in RP translation upon mGlu_1/5_ stimulation led us to wonder whether new RP synthesis was required for mGluR-LTD (**Fig. 3a**). Ribogenesis requires the coordinated activation of RNA polymerase (Pol) I to transcribe 45S precursor rRNA, RNA Pol II to transcribe RP transcripts, and RNA Pol III to transcribe 5S rRNA ^40^. To block the translation of RPs, we chose the selective Pol I inhibitor CX-5461, which blocks transcription of rRNA and the subsequent transcription of RPs without disrupting general transcription mediated by RNA Pol II ^41,42^ (**Fig. 3a**). Previous work has shown CX-5461 disrupts ribogenesis in acute hippocampal slices while preserving overall cell health and baseline activity^43^. To confirm the efficacy of CX-5461 in our system, we incubated hippocampal slices with 200 nM drug or vehicle and measured nucleolar volume in CA1 pyr neurons using fibrillarin immunostaining followed by confocal imaging and 3D reconstruction. We find that nucleolar volume is significantly reduced in both WT and *Fmr1^-/y^* hippocampal slices within 45 min of incubation with CX-5461 (**Fig. 3b**). Next, we tested whether CX-5461 blocked the increased RP translation stimulated by mGlu_1/5_ activation. Slices were pre-incubated in 200 nM CX-5461 for 30 min and stimulated with DHPG for 5 min followed by a 25 min washout. The translation of *Rps25* was measured using TRAP and qPCR. Our results show that DHPG induces a significant increase in *Rps25* translation in WT slices that is blocked in the presence of CX-5461 (**Fig. 3c**). Consistent with the RNA-seq results, the increase in *Rps25* translation is impaired in *Fmr1^-/y^* slices, and this is not significantly altered with CX-5461.

**Figure 3.**
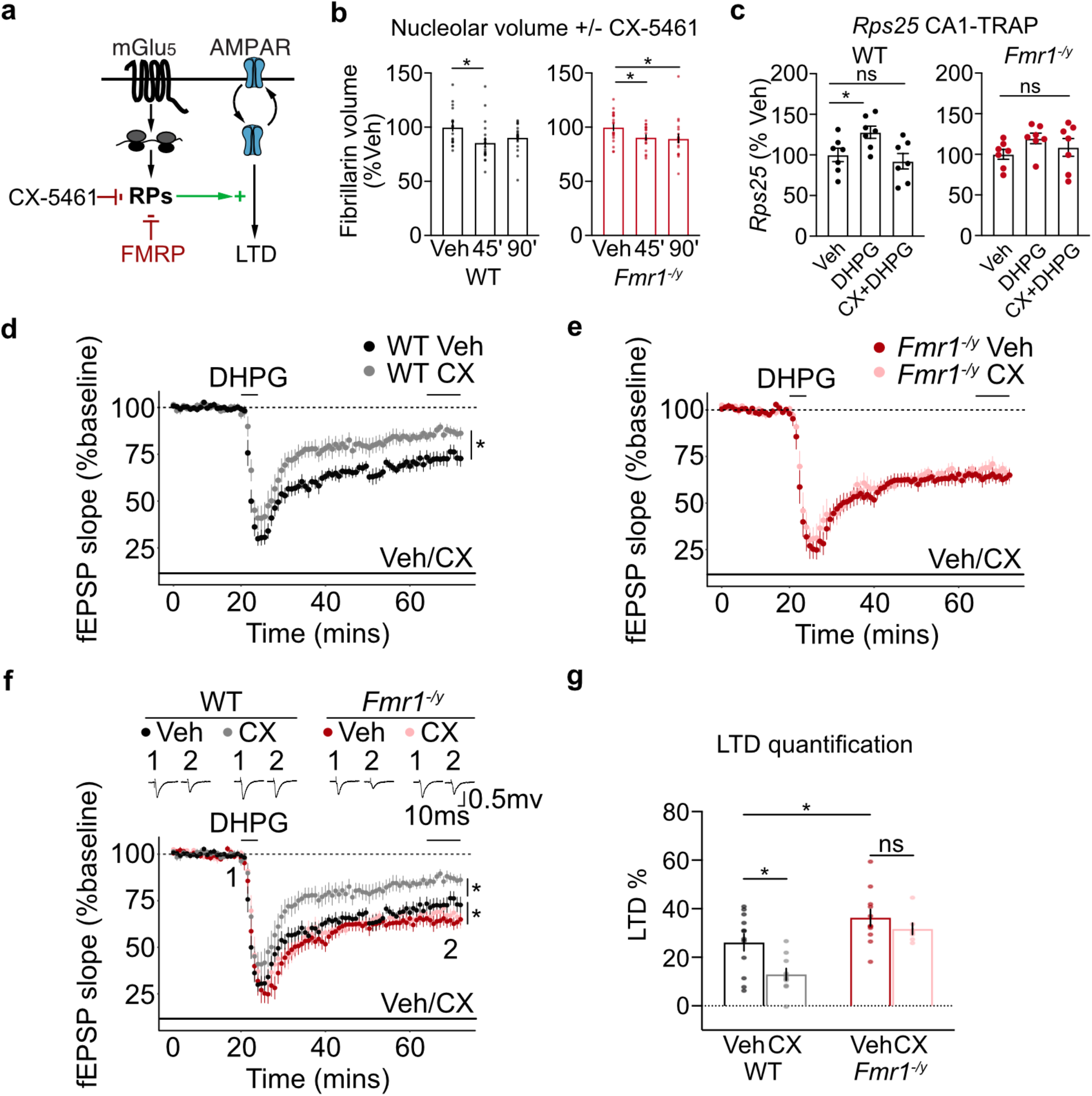
RP translation is required for mGluR-LTD in WT but not in *Fmr1^-/y^* hippocampal CA1. **(a)** Schematic of experimental strategy. Specific RNA Pol1 blocker CX-5461 was used to ribogenesis, including RP translation. **(b)** Fibrillarin staining of hippocampal slices shows that nucleolar volume of CA1 pyr neurons is significantly reduced in both genotypes with 200 nM CX-5461 treatment by 45 minutes (WT: one-way ANOVA *p = 0.0206, post-hoc Veh-45’ *FDR = 0.0141, Veh-90’ FDR = 0.070, *Fmr1^-/y^*: one-way ANOVA *p = 0.0488, post-hoc *Veh-45’ FDR = 0.0485, *Veh-90’ FDR = 0.0485, n = 20 neurons per group). **(c)** CA1-TRAP and qPCR analysis of *Rps25* translation in hippocampal slices confirms an upregulation in the WT in response to DHPG at 30’ that is blocked with 200 nM CX-5461. DHPG does not elicit a significant increase in Rps25 translation in *FmrF^-/y^* slices nor does CX-5461 have a significant effect (WT: one-way ANOVA *p= 0.0388, post-hoc Veh-DHPG *FDR = 0.0121, Veh-CXDHPG FDR = 0.3506, *Fmr1^-/y·^* one-way ANOVA p = 0.2627). **(d-e)** mGluR-LTD was measured in hippocampal CA1 in the presence of vehicle or CX-5461. Slices were prepared from *Fmr1^-/y^* and WT, recovered for at least 2 hours, and exposed to vehicle or 200 nM CX-5461 for at least 30 min prior to DHPG application and throughout the LTD recording. CX-5461 causes a significant reduction in LTD magnitude in WT CA1 versus vehicle (Veh = 26.13% ± 3.5% n=12, CX5461 = 13.00 % ± 2.4% n = 10, *FDR = 0.0031). In contrast, CX-5461 does not significantly reduce LTD magnitude in *Fmr1^-/y^* CA1 versus vehicle(Veh = 36.337% ± 3.9% n = 11, CX-5461 = 31.717% ± 2.3% n = 7, FDR = 0.1224) **(f-g)** Quantification of all 4 groups show a differential effect of CX-5461 on LTD in each genotype (Two-way ANOVA, genotype *p <0.001 treatment *p = 0.01, post-hoc: WT-Veh vs *Fmr1^-/y^*-Veh *FDR = 0.0107, WT-Veh vs WT-CX5461 *FDR = 0.0031, *Fmr1^-/y^ -Veh* vs *Fmr1^-/y^*–CX-5461 FDR = 0.1224).

Having confirmed that CX-5461 blocks the RP translation downstream of mGlu_1/5_ activation, we next assessed the impact on mGluR-LTD. Hippocampal slices were prepared from juvenile *Fmr1^-/y^* and WT littermates as in previous studies, with recordings performed blind to genotype and in an interleaved fashion. Vehicle or 200 nM CX-5461 was pre-incubated for at least 1 hr, LTD induced with a 5 min application of 50 *μ*M S-DHPG and fEPSPs were recorded over 1 hour. We find that CX-5461 significantly impairs LTD in WT slices, indicating that ribosomal protein translation is required for mGluR-LTD (**Fig. 3d**). However, in *Fmr1^-/y^* slices where ribosomes are overexpressed, CX-5461 has no impact on mGluR-LTD magnitude (**Fig. 3e**). The significant elevation in vehicle-treated WT and *Fmr1^-/y^* slices confirms the exaggerated LTD phenotype is present, and not impacted by CX-5461 (**Fig. 3f-g**). These results show that preventing the RP translation downstream of mGlu_1/5_ activation impairs mGluR-LTD in WT, but not *Fmr1^-/y^* hippocampus.

### *Fmr1^-/y^* neurons exhibit a length-dependent imbalance in mRNA translation that disfavors long transcripts

Our results show that RPs are overtranslated in *Fmr1^-/y^* neurons, translated downstream of mGlu_1/5_, and are required for LTD. However, the mechanism by which a change in ribosome availability could result in LTD remained unclear. Along with the increase in RPs, our TRAP-seq results showed that a consequence of LTD is the downregulation of synaptic transcripts, which tend to be longer than average ^44^. In contrast, we observed an upregulation of transcripts encoding metabolic proteins, which tend to be short ^45^. Interestingly, a conserved property seen in cells from yeast to humans is that the rate of protein production from shorter mRNAs is greater than that of longer transcripts^46–48^. Modelling studies have proposed that this difference could be due to the slower rate at which translation initiation occurs on long transcripts, often due to secondary structures present in the elongated 5’ untranslated region (5’UTR) ^48–50^. It has also been suggested that the smaller distance between 5’ and 3’ UTRs gives a competitive advantage to mRNAs with shorter coding sequences (CDSs) by facilitating ribosome re-initiation ^47^. These models predict a change in the concentration of ribosomes will disproportionately impact shorter mRNAs because they are more rapidly translating ^47,51,52^. This prediction is consistent with recent work showing that RP mutations that reduce cellular ribosome expression exhibit a disproportionately greater reduction in the translation of short versus long mRNAs ^42^.

Based on these studies, we wondered whether the increased ribosome availability seen in *Fmr1^-/y^* neurons and in DHPG-stimulated neurons changed the profile of translating mRNAs in a length-dependent manner. If so, this would result a reduced translation of long mRNAs encoding synaptic proteins. This change in the composition of the translating population would be consistent with a sustained depression of synaptic strength (**Fig. 4a**). This model predicts that (1) FX neurons exhibit a reduced translation of long mRNAs, (2) the long mRNAs under-translated in *Fmr1^-/y^* neurons are associated with synaptic stability, (3) stimulation of mGlu_1/5_ induces the same reduction in long mRNA translation in WT, (4) inhibiting ribosome protein translation prevents the reduction of long mRNA translation upon mGlu_1/5_ stimulation.

**Figure 4.**
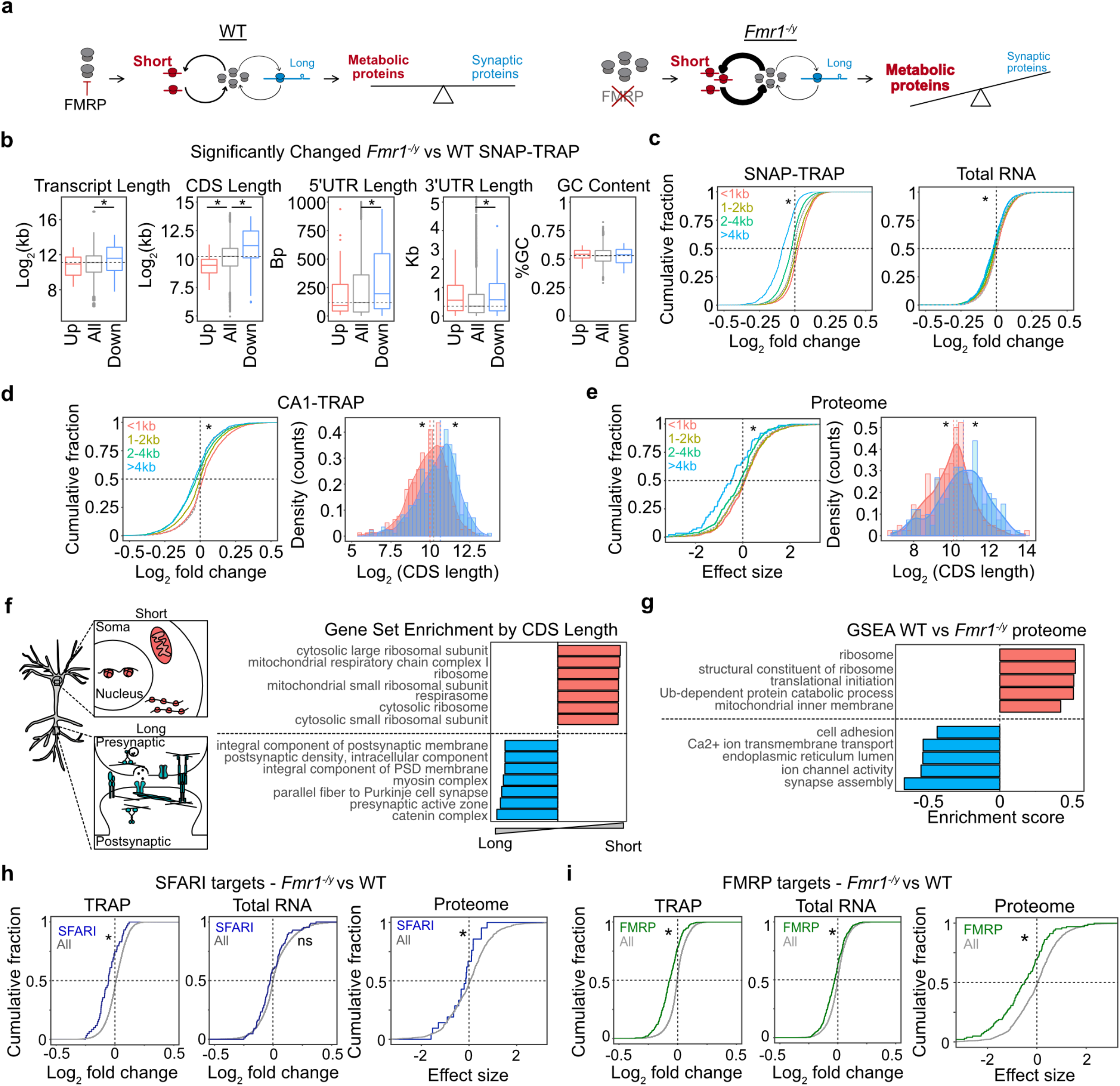
Under-translation of long mRNAs causes reduced expression of synaptic proteins, autism risk factors, and FMRP targets in *Fmr1^-/y^* hippocampal neurons. **(a)** A model of altered translation in FX proposes that increased ribosome production changes the differential rate of translation between short and long mRNAs, which ultimately alters the proportion of proteins with metabolic versus synaptic functions. **(b)** The significantly altered *Fmr1^-/y^* SNAP-TRAP population (DESeq2 adjusted p-value < 0.1) exhibits an imbalance in transcript length, CDS length, 5’UTR length and 3’UTR that indicates under-translation of long, low-initiation mRNAs (Transcript length: Kruskal Wallis *p = 4.01X10^-5^, Wilcoxon rank sum test all vs up p = 0.733, all vs down *p = 7.28X10^-6^, CDS length: Kruskal Wallis *p = 2.219X10^-14^, Wilcoxon rank sum test all vs up *p = 3.181X10^-5^, all vs down *p = 1.516X10^-11^, 5’UTR length: Kruskal Wallis *p= 0.01179, Wilcoxon rank sum test all vs up p = 0.9794, all vs down *p = 0.00282, 3’UTR length: Kruskal Wallis *p= 0.0022, Wilcoxon rank sum test all vs up p = 0.1103, all vs down *p = 0.001827, coding GC content: Kruskal Wallis p= 0.5817). **(c)** A binned analysis of the total *Fmr1^-/y^* SNAP-TRAP population shows that the CDS length imbalance is seen in the whole ribosome bound population, with the shortest group showing the most positive log2 fold change distribution and the longest group showing the lowest log2 fold change (KS test, <1kb vs >4kb *p < 2.2X10^-16^ d = 0.537). The total transcriptome also demonstrates this property but at a much lower magnitude (KS test <1kb vs >4kb *p = 2.00X10 ^-11^ d = 0.1470). **(d)** The *Fmr1^-/y^* CA1-TRAP population shows a similar CDS length bias that can be seen in a binned analysis or a comparison of the top 500 most up- and downregulated mRNAs ranked by log_2_ fold change (KS test <1kb vs >4kb *p < 2.2X10^-16^, two sample z test all vs up z = −3.416, *p = 0.0006356, all vs down z = 5.0454, *p = 4.525 ×10^-7^). **(e)** The *Fmr1^-/y^* proteome exhibits a CDS length-dependent imbalance that mirrors the results seen by TRAP, both in a binned analysis (KS test <1kb vs >4kb *p = 2.84X10^-6^) and comparison between the 200 most up- and downregulated proteins (two-sample z test, up vs all: z = −2.8216 *p = 0.005, down vs all: z = 3.0408, *p = 0.0024). **(f)** A Gene set analysis ranked by CDS length reveals that shorter transcripts are enriched with ribosome and mitochondrial genes whereas longer transcripts are enriched with synaptic structure related genes. **(g)** GSEA of the *Fmr1^-/y^* proteome shows a significant downregulation of synaptic gene sets and a significant upregulation of ribosome/mitochondrial gene sets. **(h)** SFARI targets are downregulated in the *Fmr1^-/y^* SNAP-TRAP population (two sample z test, z = −5.43 p = 5.65X10^-8^) and not changed in the total RNA population (two sample z test, z = −1.64, p = 0.099). SFARI targets are significantly underexpressed in the *Fmr1^-/y^* proteome, consistent with the under-translation seen by TRAP (two sample z test, z = −2.0008, *p = 0.045). **(i)** FMRP targets are significantly downregulated in the *Fmr1^-/y^* SNAP-TRAP population (two sample z test, z = −12.461 *p < 2.2X10^-16^) and changed to a lesser degree in total RNA population (two sample z test, z = −5.42, p = 5.805X10^-8^). FMRP targets are significantly reduced in the *Fmr1^-/y^* proteome (two sample z test, z = 5.30, *p = 1.135 × 10^-7^).

To test the first prediction, we investigated whether *Fmr1^-/y^* neurons exhibit a length-dependent change in the translating mRNA population by comparing the significantly altered population in the *Fmr1^-/y^* SNAP-TRAP fraction. This analysis revealed a striking relationship between differential expression and transcript length in the *Fmr1^-/y^* TRAP. The significantly downregulated population contains transcripts with longer CDS regions versus the average population, and the upregulated population contains transcripts with shorter CDS regions (**Fig. 4b**). Downregulated mRNAs also contain significantly longer 5’ UTRs, consistent with a reduced initiation rate. An alteration in translation GC content of the CDS, which has also been implicated in translation efficiency, is not associated with differential expression ^50^. We next examined differential expression in the total population by comparing groups binned by CDS length < 1 kb, 1-2 kb, 2-4 kb and > 4 kb. This analysis again shows a significant shift in *Fmr1^-/y^* TRAP fraction that disfavors long transcripts (**Fig. 4c**). The same imbalance is seen in the total transcriptome, however the magnitude of this change is significantly smaller. This indicates that the effect seen in the TRAP is not driven by a change in transcript abundance.

Our next question was whether the same length-dependent shift is seen *Fmr1^-/y^* CA1 pyr neurons, so we performed the same analysis on the CA1-TRAP population. Our results show the same relationship between fold change and CDS length seen in the SNAP-TRAP population **(Fig. 4d**, **Supplementary Fig. 3a**). To ensure this difference was not due to a difference in transcript number between groups, we performed an additional analysis comparing an equal number of transcripts (500) in the up- and down-regulated population when ranked by nominal p-value (**Fig. 4e**). This confirmed a significant difference in length between up- and downregulated transcripts in the *Fmr1^-/y^* CA1-TRAP population. This imbalance persists when normalized to the total transcriptome, and is not due to a systemic length bias in the RNA-seq dataset (**Supplementary Fig. 3b-c**)^53^. Together these results indicate an under-translation of long mRNAs in *Fmr1^-/y^* neurons.

Interestingly, the length of RP transcripts is significantly shorter than average, raising the possibility that the upregulation of RPs is a be a consequence rather than the cause of the length-dependent shift. However, further analyses show that the RP population is significantly more upregulated than equal-length transcripts, suggesting it is not driven by the elevated translation of short mRNAs in *Fmr1^-/y^* neurons (**Supplementary Fig. 3d**).

### The length-dependent translation shift in *Fmr1^-/y^* neurons reduces expression of synaptic proteins, autism risk factors, and FMRP targets

Given the length-dependent imbalance in translation seen in *Fmr1^-/y^* neurons, we next asked whether this was mirrored in a length-dependent change in protein expression. We therefore performed an analysis of length on the most significantly up and down-regulated proteins in the *Fmr1^-/y^* hippocampal proteome (**Fig. 4e**). Consistent with our TRAP-seq results, we find the population downregulated in the *Fmr1^-/y^* is enriched for proteins encoded by longer CDS transcripts, with proteins encoded by transcripts > 4kb exhibiting the most significant downregulation (**Fig. 4e**). Similar to TRAP-seq, there is no correlation between significance and protein length (**Supplementary Fig. 3e**).

The second prediction of our model is that the long mRNAs under-translated in *Fmr1^-/y^* neurons encode proteins involved in synaptic stability. To determine whether mRNA length was a factor in functional segregation in the CA1 translating mRNA population we performed a gene set analysis ranked by CDS length (**Fig. 4f, Table 8**). This reveals longer CDS transcripts are enriched for synaptic adhesion molecules, pre- and post-synaptic assembly proteins, and calcium channels, while shorter transcripts are enriched for nuclear, ribosomal, and mitochondrial proteins. The CDS length-dependent imbalance in translation in FX neurons therefore predicts an underproduction of proteins important for synaptic function. To test this, we employed GSEA on the *Fmr1^-/y^* proteome and confirm that the most significantly enriched categories in the underexpressed population are related to synaptic structure and function (**Fig. 4g, Supplementary Fig. 3f**). Although differences in biological fractionation and analysis make it difficult to compare significant differences in individual targets identified in the *Fmr1^-/y^* proteome to those identified by TRAP-seq, we nevertheless find there is a significant overlap between the populations of targets significantly downregulated in the *Fmr1^-/y^* versus WT (p < 0.05) (**Supplementary Fig. 3).** Overlapping targets include proteins involved in synaptic structure that have previously been implicated in FX including Shank1 and Map1b. **(Supplementary Tables 9-10**). These results show that the long synaptic mRNAs downregulated in the *Fmr1^-/y^* TRAP are consistent with those reduced in the *Fmr1^-/y^* proteome.

Interestingly, it has been noted that autism risk factors identified by the Simons Foundation Autism Research Initiative (SFARI) are encoded by long genes^54–56^. A downregulation of these targets would be predicted to have deleterious consequences on neuronal function, and we therefore investigated whether this group was reduced in *Fmr1^-/y^* neurons. Our results show that SFARI transcripts are significantly downregulated in the *Fmr1^-/y^* SNAP-TRAP and CA1-TRAP populations, which is consistent with the increased length of this population in our dataset (**Fig. 4h, Supplementary Fig. 3h**). Furthermore, there is a significant correlation between differential expression and length even within the SFARI population (**Supplementary Fig. 3i**). The same downregulation is not seen in the total *Fmr1^-/y^* transcriptome, indicating it is not driven by reduced abundance (**Fig. 4h, Supplementary Fig. 3i)**. Additional analyses show that the length of the CDS, and not the gene, is driving reduced expression in the TRAP population (**Supplementary Fig. 3j-k**). Importantly, examination of the *Fmr1^-/y^* proteome reveals an underexpression of proteins encoded by SFARI targets, indicating the change in translation has an impact on the steady-state proteome (**Fig. 4h**).

FMRP targets are also longer in length versus other neuronal transcripts and we and others have shown these targets are underexpressed in the ribosome-bound fraction of *Fmr1^-/y^* CA1 neurons ^21,57,58^. Our results confirm that FMRP targets are reduced in the *Fmr1^-/y^* SNAP-TRAP fraction consistent with results from CA1 neurons, with a small but significant effect seen in the total transcriptome (**Fig. 4i**). However, it was not clear that this downregulation would indicate a reduction in protein expression, as recent studies indicate the reduced ribosome binding of these targets may represent accelerated elongation ^11,59^. To investigate this question, we quantified FMRP targets in the *Fmr1^-/y^* proteome, and found the same downregulation seen in the TRAP fraction (**Fig. 4i**). Together, these results confirm an underexpression of SFARI risk factors and FMRP targets that is consistent with the reduced translation of long transcripts.

### mGluR-LTD mimics the length-dependent translation imbalance seen in *Fmr1^-/y^* neurons

According to the third prediction of our model, the reduced translation of long mRNAs should be a consequence of mGluR-LTD, and this should be occluded in *Fmr1^-/y^* neurons (**Fig. 5a**). To test this, we performed the same analysis of transcript length in the DHPG-induced CA1-TRAP fraction. Remarkably, we find that the same decrease in long mRNAs observed in *Fmr1^-/y^* CA1 neurons is induced in WT neurons with application of DHPG (**Fig. 5b**). Similar to the downregulated population in *Fmr1^-/y^* neurons, these transcripts exhibit a significantly longer 5’UTR indicative of a reduced initiation rate. Analysis of the total significantly altered population binned by kb shows that upregulated targets are significantly shorter than average, and downregulation targets are significantly longer than average, effects that are not seen in the total transcriptome (**Fig. 5c, Supplementary Fig. 4a, Supplementary Table 11**). Next, we investigated whether the basal length-dependent shift in the basal *Fmr1^-/y^* CA1-TRAP population impairs a further shift with DHPG. Our analysis of the *Fmr1^-/y^* CA1-TRAP population shows that there is a small but significant shift in length in the CA1-TRAP after DHPG stimulation, however this is much smaller than that observed in WT (**Supplementary Fig. 4b**). To more directly compare the impact of DHPG between genotypes, we examined the 500 most up- and downregulated transcripts. This analysis reveals a significant divergence in length between the up- and down-regulated populations in DHPG-treated WT slices that is significantly reduced in DHPG-treated *Fmr1^-/y^* slices (**Fig. 5d**). These changes are not seen in the total transcriptome (**Supplementary Fig. 4c**). These results show that the length-dependent shift seen with DHPG stimulation is occluded in the *Fmr1^-/y^* hippocampus.

**Figure 5.**
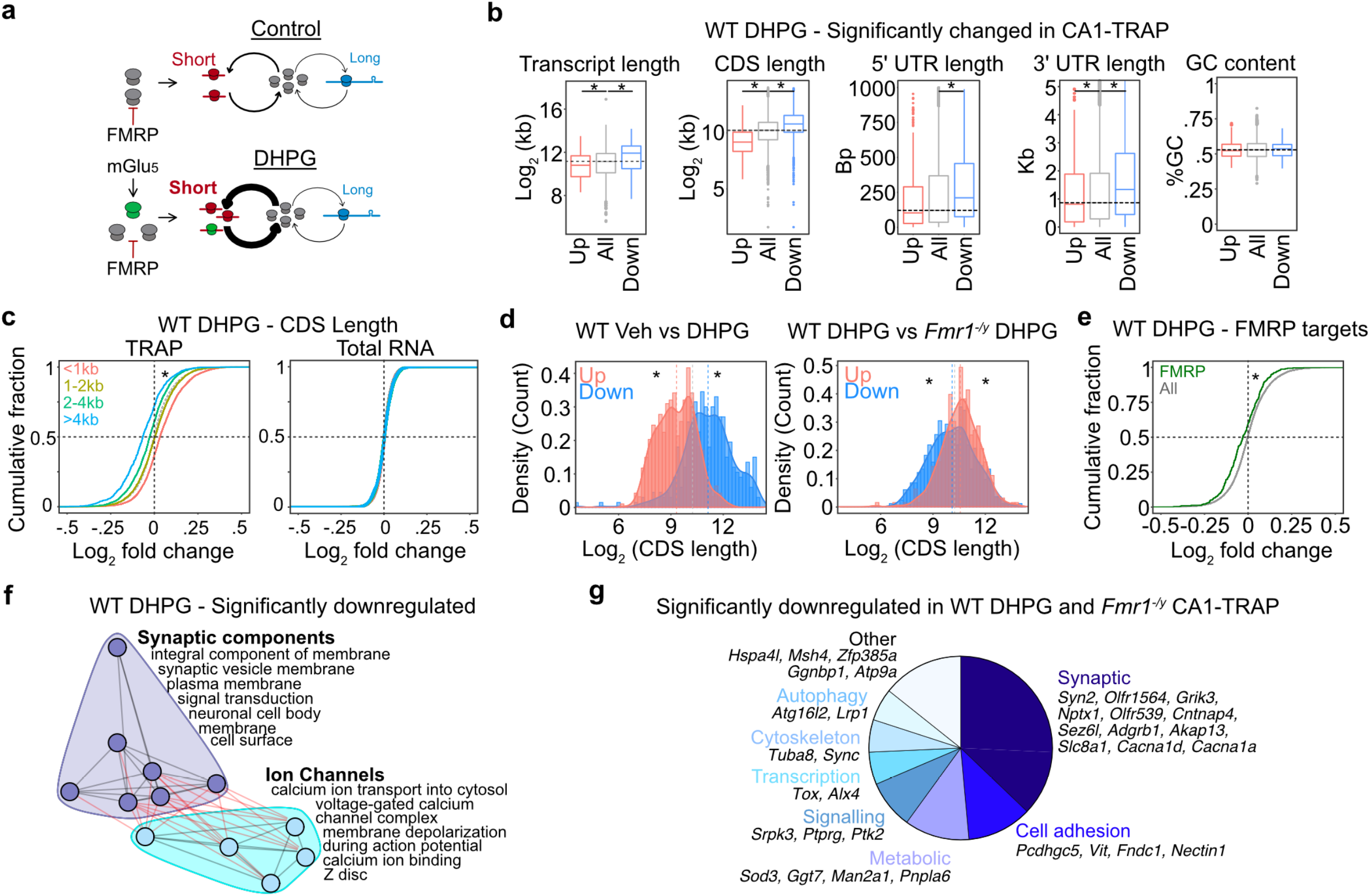
Translation of long mRNAs is reduced in CA1 pyr neurons with induction of mGluR-LTD. **(a)** Our model predicts that the increased ribosome production seen with mGluR-LTD will cause a similar length-dependent imbalance as seen in the *Fmr1^-/y^* translating population. **(b)** Analysis of the significantly changed population in WT DHPG CA1-TRAP shows a significant imbalance in transcript length, CDS length, 5’UTR length, and 3’UTR length that matches the basal *Fmr1^-/y^* CA1-TRAP population (Transcript length: Kruskal Wallis p = 2.105X10^-14^, Wilcoxon rank sum test all vs up p = 6.042X10^-5^, all vs down p = 1.022X10^-11^, CDS length: Kruskal Wallis *p < 2.2X10^-16^, Wilcoxon rank sum test all vs up *p < 2.2X10 ^-16^, all vs down *p < 2.2X10^-16^, 5’UTR length: Kruskal Wallis *p= 0.0005, Wilcoxon rank sum test all vs up p = 0.0808, all vs down *p = 0.0006, 3’UTR length: Kruskal Wallis *p= 0.000, Wilcoxon rank sum test all vs up *p = 0.03821, all vs down *p = 0.0002, coding GC content: Kruskal Wallis p= 0.9375). **(c)** A binned analysis shows that there is a CDS length-shift in the TRAP fraction (KS test <1kb vs >4kb *p < 2.2X10^-16^) and no change in the total transcriptome (KS test <1kb vs >4kb p = 0.698). **(d)** Comparison of the top and bottom 500 differentially expressed transcripts in WT DHPG shows a significant effect of length (All vs up z = −17.831, *p < 2.2X10^-16^, All vs Down z = 10.774, *p < 2.2X10^-16^). Comparison between WT DHPG vs *Fmr1^-/y^* DHPG reveals that the length shift is occluded in *Fmr1^-/y^* (all vs Up z = 5.2982, *p = 1.16 X10^-7^, all vs Down z = −2.2827, *p =0.02). **(e)** As predicted by their longer lengths, FMRP targets are reduced with DHPG in the WT CA1-TRAP population (two sample z test, z = 5.333, p = 9.66 x 10^-8^). This change is not seen in the total transcriptome (two sample z test, z = −2.039, p = 0.041). **(f)** Network analysis of the downregulated GO terms reveals a concentration of synaptic components and ion channel clusters. **(g)** Analysis of the population significantly downregulated in both WT DHPG and *Fmr1^-/y^* CA1-TRAP fractions (p < 0.05) identifies 42 transcripts, many of which are involved in synaptic function.

We next asked whether there was a functional similarity in the transcripts downregulated with DHPG in the WT TRAP versus those basally reduced in the *Fmr1^-/y^* TRAP. First, we asked whether the reduction in long mRNAs coincides with a reduction in FMRP targets, similar to what is seen in the *Fmr1^-/y^* TRAP. Our results show that there is a significant reduction in FMRP targets in the WT CA1-TRAP after DHPG stimulation, which is not seen in the total transcriptome (**Fig. 5e, Supplementary Fig. 4d**). Next, we performed a clustering analysis of GO terms enriched in the population downregulated in WT DHPG. This revealed that downregulated targets largely fell into categories related to synaptic components and ion channels, similar to what is seen in the *Fmr1^-/y^* proteome (**Fig. 5f**). Downregulated targets include those related to synaptic signalling and structural stability including cadherins/protocadherins (*Pcdhac2, Pcdh1, Celsr3, Celsr2, Cdh18, Cdh2, Pcdhgc5*, etc.) and cell adhesion molecules (*L1cam, Nrcam, Focad, Cadm3*, etc.) (**Supplementary Fig. 4e**). In addition, the downregulated population contains multiple targets that are involved in calcium regulation downstream of mGlu_1/5_ activation, including voltage-gated calcium channel transcripts (*Cacna1b, Cacna1c, Cacna1i, Cacna1ad2*) and ryanodine receptors (*Ryr2 and Ryr3*) ^60^ (**Supplementary Fig. 4f**). To determine the similarity between the specific transcripts altered with DHPG in WT and those basally altered in the *Fmr1^-/y^* CA1-TRAP population, we compared the targets significantly downregulated in both populations (p < 0.05). This analysis shows a significant overlap, with the majority of the 42 overlapping targets related to synaptic function (**Fig. 2g, Supplementary Fig. 4g, Supplementary Table 12**). Together, these results show that transcripts downregulated with DHPG in WT are functionally similar to those basally downregulated in the *Fmr1^-/y^* CA1-TRAP.

### Inhibition of ribosomal protein production prevents mGluR-induced changes in translation

A critical test of our model is whether disruption of ribosomal protein translation alters the length-dependent change in translation downstream of mGluR-LTD (**Fig. 6a**). To test this fourth prediction, we stimulated hippocampal slices with DHPG to induce LTD in the presence of 200 nM CX-5461 and performed TRAP-seq to identify CA1 neuron specific translation (**Supplementary Fig. 5a, Supplementary Table 13**). Consistent with our first TRAP-seq experiment, we find that DHPG stimulates RP translation in WT neurons, and this is significantly impaired in *Fmr1^-/y^* neurons (**Supplementary Fig. 5b**). In stark contrast, incubation with CX-5461 blocks the increase in RP translation stimulated by DHPG in WT neurons and results in a small but significant downregulation in RPs in *Fmr1^-/y^* slices that may be due to elevation in the basal population (**Fig. 6b**). Remarkably, analysis of the 500 most up- and downregulated targets reveals that CX-5461 also eliminates the length-dependent shift in translation induced by DHPG (**Fig. 6c**). This provides a causal link between RP translation and the length-dependent shift in translation induced by mGlu_1/5_ activation. Importantly, CX-5461 does not alter the upregulation of immediate early genes such as *Npas4* with DHPG in either genotype, indicating there is no impairment of the general responsiveness to mGlu_1/5_ stimulation (**Supplementary Fig. 5c-d, Supplementary Table 13**).

**Figure 6.**
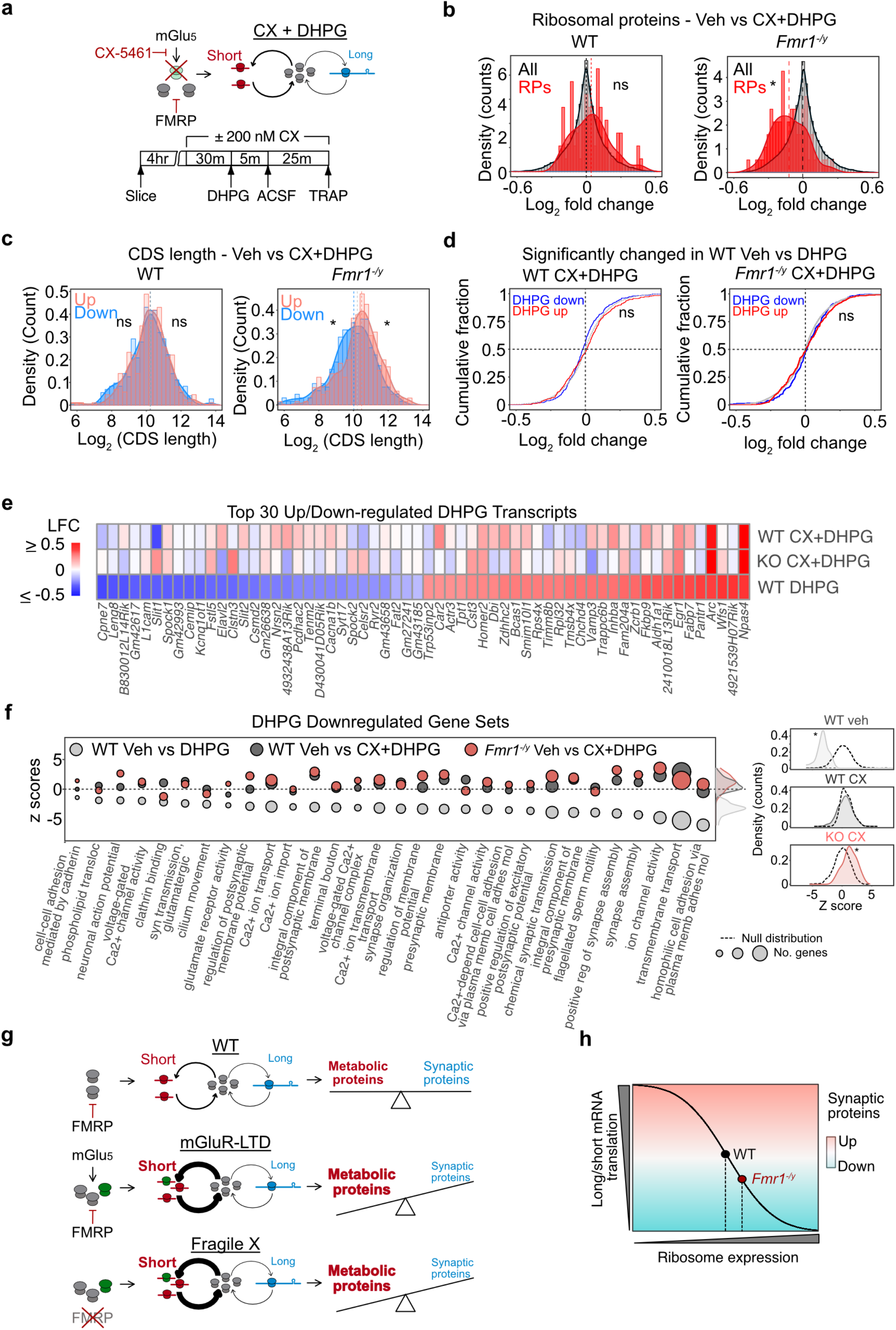
Inhibition of RP translation prevents reduction of long mRNA translation during mGluR-LTD. **(a)** Our model predicts that preventing the increase in RP translation downstream of mGlu_1/5_ activation should prevent the length-dependent shift in translation. Experimental timeline for CX+DHPG CA1 TRAP-seq experiment is shown. (**b**) Incubation with CX-5461 blocks the increase in RP translation seen with DHPG stimulation in the WT CA1-TRAP population (RP: two sample z test, z = 0.79134, p = 0.428741). CX-5461 causes a slight reduction in RP translation with DHPG in *Fmr1^-/y^* CA1-TRAP (RP: two sample z test, z = −4.30, *p = 1.69 × 10^-5^). (**c**) The length-dependent shift in translation seen with DHPG in WT is eliminated with incubation of CX-5461 (Top 500: two sample z test, All vs up z = 0.2897, p 0.77200, All vs down z = 0.10510, p 0.916292). A slight reversal of the length-dependent imbalance evoked by DHPG is also seen in the *Fmr1^-/y^* CA1-TRAP (Top 500: two sample z test, All vs Up z = 3.1899, *p = 0.0014, All vs down z = −2.282, *p = 0.022). (**d**) LTD transcripts were defined as significantly regulated the first in WT DHPG TRAP-Seq dataset. Incubation with CX-5461 eliminates the up- and downregulation of LTD transcripts with DHPG application in WT (two sample z test, LTD up: z = 1.619, p = 0.1054, LTD down: z = −0.9977, p = 0.3184). Application of CX-5461 has no impact on the response of LTD transcripts to DHPG in the *Fmr1^-/y^* CA1-TRAP (two sample z test up z = 1.838, p = 0.0659; down z = 1.694, p = 0.0901). **(e)** A heatmap of log2 fold change shows the significant impact of CX-5461 on the 30 most up- and downregulated LTD transcripts in WT and *Fmr1^-/y^* CA1-TRAP. The preserved upregulation of immediate early genes *Npas4* and *Arc* is highlighted. **(f)** Gene sets significantly downregulated with LTD in WT are no longer downregulated with CX-5461 in either genotype. Z tests of the distribution of LTD downregulated gene sets show that significant downregulation in vehicle-treated WT (two sample z test, z = −18.24, *p <2.2X10^-16^) is no longer changed after incubation with CX-5461 (two sample z test, WT CXDHPG: z = −0.15442, p = 0.8773), In the *Fmr1^-/y^* CA1-TRAP, CX-5461 causes a slight upregulation of these gene sets, indicating a reversal in the translation regulation of these mRNAs (two sample z test, z = 4.4808 *p = 7.435X10^-6^) **(g)** Our results fit a model whereby activation of mGlu_1/5_ causes an increase in ribosome production that drives an imbalance in the translation of short versus long mRNAs. This imbalance is similarly driven by the loss of FMRP, which increases ribosome production and mimics the LTD translation state. Ultimately, the reduced translation of long mRNAs reduces expression of proteins that participate in synaptic stability and function. **(h)** We propose that the altered translation imbalance driven by excessive ribosome production may underlie a number of phenotypes in FX that are derived from synaptic weakening or instability.

To determine whether CX-5461 had a significant impact on the translation of DHPG-regulated targets, we analysed the expression of the transcripts identified in our first DHPG TRAP-seq experiment. Validating our original results, targets significantly up- and downregulated in our first dataset are significantly up- and downregulated in the Veh WT DHPG population of the CX+DHPG dataset (**Supplementary Fig. 5e**). Consistent with the impairment of mGluR-LTD, application of CX-5461 eliminates the differential expression of translating mRNAs seen with DHPG stimulation (**Fig. 6d**). Examination of individual DHPG-stimulated targets shows that CX-5461 prevents the differential translation of most, with the exception of immediate early genes such as *Npas4, Arc*, and *Junb*, which are transcribed in response to changes in neuronal activity (**Fig. 6e**). To assess the functional impact of CX-5461 on the LTD response, we examined the gene sets identified by GSEA in the population of mRNAs normally downregulated in the DHPG CA1-TRAP. Our results show that these gene sets are no longer altered in the presence of CX-5461 in WT, and are mildly reversed in the *Fmr1^-/y^* population. The downregulated transcripts most impacted by CX-5461 are related to cell adhesion, ion channel activity, and synapse assembly (**Fig. 6f**). These results show that inhibition of RP translation impairs a translation profile that supports synaptic weakening upon stimulation with DHPG. Together, our results suggest that an imbalance in the translation of longer versus shorter mRNAs is induced by RP translation downstream of mGlu_1/5_ activation, and this translation state is basally exacerbated in *Fmr1^-/y^* CA1 neurons (**Fig. 6g**).

## Discussion

This study provides novel insights into the role of protein synthesis in FX. First, it identifies ribosomal protein transcripts as the major gene category over-translated in *Fmr1^-/y^* hippocampal neurons, which is manifest as an overexpression of ribosomes. This elevation in RP translation coincides with a length-dependent shift in the translating mRNA population that favors shorter versus longer mRNAs, a profile that reduces the expression of synaptic proteins, SFARI risk factors and FMRP targets. Second, this first cell type-specific translation profiling of mGluR-LTD shows that stimulation of mGlu_1/5_ causes a robust translation of RPs in CA1 neurons that is required for LTD. This effect is mimicked and occluded in the *Fmr1^-/y^* hippocampus. Remarkably, the increase in RP translation coincides with a reduction in the translation of long transcripts encoding synaptic proteins, and this can be blocked with ribogenesis inhibitor CX-5461. This suggests mGlu_1/5_ activation induces a length-dependent alteration in the translating population of CA1 pyr neurons that facilitates LTD through a reduced production of synaptic elements (**Fig. 6h**).

Our results are consistent with previous work showing that changes in ribosome expression alter the composition of the translating mRNA population in a length-dependent manner^42,51^. By increasing ribosome availability, we suggest that the translation of shorter versus longer mRNAs has become exacerbated in *Fmr1^-/y^* neurons due to the competitive advantage of short transcripts with a higher translation efficiency. We show that the paradoxical reduction in the translation of long mRNAs in *Fmr1^-/y^* neurons mimics the translation state of LTD and coincides with a reduced expression of proteins involved in synaptic function, many of which are implicated in autism. Although the effect sizes seen in our RNA-seq and proteomics datasets are relatively small, the modest magnitude of these changes is consistent with the hallmark *Fmr1^-/y^* phenotypes of exaggerated mGluR-LTD and excessive protein synthesis, which are expressed no more than 15-20% above WT ^4,13^. These phenotypes have provided important direction for both understanding the biology of FX and for identifying therapeutic strategies including mGlu_5_ antagonists and lovastatin ^18,61,62^. The prediction from this current study is that a long-term reduction in ribogenesis would alter the translating population in *Fmr1^-/y^* neurons to resemble WT, thereby increasing the production of synaptic elements. Future work is needed to test this hypothesis.

A key question raised by our study is whether the elevation in ribogenesis we observe is a primary or secondary consequence of FMRP loss. FMRP binding partners encode multiple signaling molecules, including regulators of the extracellular-regulated kinase (ERK) and mammalian target of rapamycin (mTOR) signaling cascades that regulate ribogenesis^3^. Although RP transcripts have not been identified as stringent FMRP binding partners, several have been identified as “low-binding” targets in recent work^58^. It is possible that the dysregulated stability or translation of these targets in the absence of FMRP results in an increased ribosome expression. As FMRP has been implicated in directly binding the 80S ribosome, it is also possible that a change in structural stability contributes to the altered ribosome expression^63^. To completely address this question, it will be necessary to investigate the impact of proximal FMRP loss and re-expression on ribogenesis in adult neurons.

Several studies have shown that RP transcripts are abundant in axons and dendrites, and that RPs are locally synthesized ^35–38^. Shigeoka et al., perform a series of elegant experiments in *Xenopus* reticular ganglion neurons to show that RPs are locally translated in axons, and the newly-synthesized RPs are incorporated into existing ribosomes ^39^. Both the synthesis and incorporation of new RPs into ribosomes is elevated with the growth factor netrin-1, which induces protein synthesis to promote axon branching. Importantly, preventing the new synthesis of Rps4x reduces protein synthesis and inhibits axon branching, showing that new RP synthesis is critical for this process. More recently, work by Fusco et al. shows that RPs are translated in the dendrites of mammalian neurons and these new RPs are incorporated into existing ribosomes in response to changes in the cellular environment^35^. The question of why new RPs would be needed to increase the functional capacity of ribosomes is intriguing given the abundance of existing RPs. Both Shigeoka et al. and Fusco et al. propose that the exchange of old or damaged RPs with new ones allows for the restoration of ribosome function without creating new ribosomes, which is energetically expensive. In support of this, both studies show that the RPs with the greatest rate of exchange are those at the outer cytosolic face of the ribosome, which are the most prone to oxidative stress. Our results confirm that these RPs are upregulated with DHPG as well (**Supplementary Table 7**).

Application of cycloheximide or inhibitors of translation control signaling pathways can block mGluR-LTD when applied for 30 min-1 hour before stimulation, indicating that the proteins supporting LTD are rapidly produced ^32,64^. The rapid action of newly-synthesized RPs may seem incongruent with the rate of ribosome turnover, which is measured in days, however RP exchange on existing ribosomes can impact translational capacity on a shorter timescale ^65^. In dendrites, the translation of RP transcripts visualized with Puro-PLA labelling in as little as 5 minutes, and newly-translated RPs can quantified in mature ribosomes within 1-2 hours. This timescale is consistent with our results showing RP transcripts are elevated in the TRAP fraction of CA1 neurons by 30 minutes post DHPG, and that inhibiting RP production has an impact on LTD at 1 hour. The reduced translation of synaptic proteins is likely due to a combination of both lower synthesis and a secondary impact on turnover. Consistent with this idea, recent work shows that changes in synaptic activity significantly alter the proteome within 2 hours through combined alterations in protein synthesis and breakdown ^66^.

Our results provide the first picture of the changing translation landscape in CA1 pyr neurons during LTD. A number of the targets altered suggest interesting mechanisms by which LTD is maintained. We find a significant upregulation of immediate early genes associated with induction of synaptic plasticity and LTD, including *Arc, Npas4* and *Egr1* ^33,34,67,68^. Regulators of signaling cascades activated by group 1 mGluRs are also upregulated in the CA1 TRAP population including MAP kinase (*mapkak3, mapk4*) and Cyclin-dependent kinase 5 (*cdk5r2*) ^69,70^. Conversely, the reduced expression of proteins involved in synaptic signaling and structure such as cadherins (*Cdh2, Cdh18, Celsr2, Celsr3*) and protocadherins (*Pcdh1, Pcdhac2, Pcdhgc5*) is consistent with a reduction in synaptic strength ^71–73^. Surprisingly, we find that several members of the Semaphorin-Plexin-Neuropilin signalling pathway (*Nrp1, Plxna2* and 3), are downregulated with LTD, suggesting that decreased semaphorin signalling might play a role in the prolonged downregulation of synaptic strength ^74^. Additionally, transcripts encoding L-type calcium channel Cav1.2 and ryanodine receptors Ryr2 and Ryr3 are downregulated with DHPG in our CA1-TRAP-seq. These targets are involved in the calcium signalling that occurs in response to mGlu_1/5_ activation ^60^, and it is possible that the downregulated expression of these proteins is involved in the saturation of further LTD seen after DHPG application ^32^. Further investigation of these and other changes seen in by CA1 TRAP-seq will be interesting for understanding the mechanisms of mGluR-LTD.

Similar to our results, previous work by Sawicka et al. shows a reduction in the expression of long FMRP targets in the TRAP fraction of *Fmr1^-/y^* CA1 neurons isolated using a Ribotag system^58^. However, the authors interpret these results as changes in transcript abundance rather than translation, reasoning that mRNAs not bound by ribosomes are quickly degraded. This is not inconsistent with our results, as we do see a small but significant reduction in long mRNAs including FMRP targets in the transcriptome of *Fmr1^-/y^* hippocampus after filtering for CA1 pyr neuron expression (**Figs 4c, 4i**). The effects seen in the CA1-TRAP fraction are much greater in magnitude and persist after normalizing to the transcriptome, which leads us to conclude that changes in this fraction are indicative of translation. However, it is important to note that we cannot rule out the possibility that underlying changes in transcript abundance partially contribute to the changes we observe as we do not quantify the transcriptome of CA1 neurons specifically. Indeed, although our study focuses on the question of translation, recent ribosome profiling studies of *Fmr1^-/y^* brain have reported changes in the transcriptome including a reduction in the stability of long mRNAs and alternative splicing of SFARI targets the *Fmr1^-/y^* transcriptome^12,75^. Although the contribution of these changes to synaptic or circuit function was not explored in these studies, the evolving picture suggests that there are multiple alterations in transcription, translation and RNA stability that ultimately contribute to neurological phenotypes in the *Fmr1^-/y^* brain.

Interestingly, a previous study in *Drosophila* shows that *Fmr1* depletion results in a reduction in the translation of long mRNAs and expression of long proteins^76^. Based on these results, the authors propose that FMRP is in fact an activator of translation for long mRNAs. In contrast, recent studies by Das Sharma et al. and Aryal et al. suggest the reduced ribosome binding of long mRNAs in ribosome profiling from *Fmr1^-/y^* cortical lysates represents an increase in the translation of these targets due to reduction in ribosome stalling by FMRP ^11,59^. Although these studies do not include a corresponding analysis of the proteome, the prediction is an increase in the production of FMRP target proteins. Our study shows that there is a significant reduction in the expression of proteins encoded by long mRNAs including FMRP targets, which is consistent with a reduction in translation (**Fig. 4i**). However, as TRAP-seq data cannot distinguish between stalled and translating ribosomes, we cannot exclude the possibility that these mRNAs are rapidly elongated but subsequently degraded, resulting in a reduction in the steady-state proteome. What we can conclude is that the reduction seen in the TRAP is indicative of the reduced expression of the encoded proteins, which is significant for determining the impact on synaptic function. Indeed, several targets we identify as reduced in translation have been implicated in FX (i.e., *Hcn2, Shank1, Pde2a*) or autism (i.e., *Cdh2, Cacna1c, Gabra5*) ^77–80^, and are consistent with the reductions in calcium channel function, GABA receptor activity and dendritic spine maturity that are seen in the *Fmr1^-/y^* mouse ^73,79,81^. These results suggest that the synaptic pathologies caused by altered translation in FX are not due to an overall increase in protein production, but rather a shift in the composition of the translating mRNA pool that favors production of metabolic proteins at the expense of longer synaptic transcripts. If so, the role of altered protein synthesis in the neuropathology of FX should not be viewed as a general over-production, but rather an imbalance that is driven by excessive ribogenesis (**Fig. 6h**). This would suggest a new conceptual framework for interpreting the role of altered translation in the development of synaptic pathology in FX.

## Methods

### Animals

*Fmr1^-/y^* and CA1-TRAP mice (created by gensat.org/ and obtained from Jackson Labs with permission from Nathanial Heintz) were bred on the JAX C57BL/6J background. *Snap25-EGFP/Rpl10a* mice were obtained from Jackson Labs and maintained on a C57BL/6J background. *Fmr1^-/y^* and WT littermates were bred using *Fmr1^+/-^* females and JAX C57BL/6J males. *Fmr1^-/y^*-TRAP and WT-TRAP littermates were bred using *Fmr1^+/-^* females and CA1-TRAP homozygous males. All experiments were carried out using male littermate mice aged P25-32 with the experimenter blind to genotype. Mice were group housed (6 maximum) in conventional non-environmentally enriched cages with unrestricted food and water access and a 12h light-dark cycle. Room temperature was maintained at 21 ± 2°C. Animal husbandry was carried out by University of Edinburgh technical staff. All procedures were in performed in accordance with ARRIVE guidelines and the regulations set by the University of Edinburgh and the UK Animals Act 1986.

### Hippocampal slice preparation

Hippocampal slices were prepared from male littermate WT and *Fmr1^-/y^* mice (P25-32), in an interleaved fashion, with the experimenter blind to genotype as described previously ^4^. Briefly, mice were anaesthetized with isoflurane and the hippocampus was rapidly dissected in ice-cold ACSF (124 mM NaCl, 3 mM KCl, 1.25 mM NaH2PO_4_, 26 mM NaHCO_3_, 10 mM dextrose, 1 mM MgCl_2_ and 2 mM CaCl_2_, saturated with 95% O_2_ and 5% CO_2_). Slices (500 μm thick) were prepared using a Stoelting Tissue Slicer and transferred into 32.5°C ACSF (saturated with 95% O2 and 5% CO2) within 5 minutes. Slices were incubated in ACSF for 4 hours to allow for recovery of protein synthesis ^4^. For DHPG stimulation, hippocampal slices were stimulated with 50 *μ*M S-3,5-Dihydroxyphenylglycine (S-DHPG) or vehicle (ddH_2_O) for 5 minutes in ACSF, then transferred to fresh ACSF for an additional 25 minutes (for TRAP and qPCR) or 55 minutes (for synaptoneurosome preparation and WB analysis). For CX-5461 experiments, slices were pre-incubated with vehicle or 200 nM CX-5461, and DHPG stimulation carried out with vehicle or 200 nM CX-5461 present throughout.

### Proteomics

#### P2 preparation

Hippocampal slices were prepared and recovered from 5 WT and *Fmr1^-/y^* littermate pairs in an interleaved and blinded fashion exactly as previously described ^4^. Slices from two mice per genotype were pooled and homogenized in 50 mM HEPES, pH 7.4 plus 0.32 M sucrose supplemented with protease inhibitor cocktail (Roche). To isolate synapse-enriched fractions, homogenates were centrifuged at 1000 x g for 10 minutes at 4°C, and the supernatant (S1) re-centrifuged at 16,000 x g for 15 minutes at 4°C. The resulting P2 pellet fractions were processed for MS analysis.

#### Filter aided sample preparation (FASP)

In-solution digestion of proteins was done as previously described ^82^. In brief, 20 μg of each sample were mixed with 75 μL 2% SDS, 1 mM Tris (2-carboxyethyl)phosphine and incubated at 55°C for 1 hour. Cysteines were blocked by adding 0.5 μL 200 mM methyl methanethiosulfonate and incubating 15 minutes at RT. After mixing with 200 μL 8 M Urea in Tris pH 8.8, samples were transferred to Microcon-30 filter tubes (Millipore) and centrifuged at 14,000 x *g* for 15 minutes at RT. Samples were washed four times with 200 μL 8 M urea and, subsequently, four times with 200 μL 50 mM ammonium bicarbonate by centrifugation as stated above. Proteins were digested with 0.7 μg Trypsin/Lys-C Mix (MS grade, Promega) in 50 mM ammonium bicarbonate overnight at 37 °C. Peptides were recovered by centrifugation. It was pooled with an additional wash with 200 μL 50 mM ammonium bicarbonate, dried in a speed vac and stored at −20 °C until used.

#### SWATH mass spectrometry analysis

Peptides were dissolved in 7 μL 2% acetonitrile, 0.1% formic acid solution containing iRT reference (Biognosys) and analyzed by micro LC MS/MS using an Ultimate 3000 LC system (Dionex, Thermo Scientific). A 5 mm Pepmap 100 C18 column (300 μm i.d., 5 μm particle size, Dionex) and a 200 mm Alltima C18 column (100 μm i.d., 3 μm particle size) were used to trap and fractionate the peptides, respectively. Acetonitrile concentration in 0.1% formic acid was increased linearly from 5 to 18% in 88 minutes, to 25% at 98 minutes, 40% at 108 minutes and to 90% in 2 minutes, at a flow rate of 5 μL/minute. Peptides were electro-sprayed into the mass spectrometer with a micro-spray needle voltage of 5500 V. Each SWATH cycle consisted of a parent ion scan of 150 msec and 8 Da SWATH windows, with scan time of 80 msec, through 450-770 m/z mass range. The collision energy for each window was calculated for a 2+ ion centered upon the window (spread of 15 eV).

The data was analyzed using Spectronaut with a spectral library previously generated from synaptosome preparations by data-dependent acquisition ^83^. The cross-run normalization based on total peak areas was enabled and the peptide abundances were exported for further processing using the R language. Only peptides quantified in at least one group with high-confidence were used, i.e., a Q-value ≤ 10^-2^ (allowing one outlier within each group). Limma R package version 3.40.6 (Bioconductor) was used to Loess normalize protein abundance (‘normalizeCyclicLoess’ function) and calculate empirical Bayes moderated t-statistics (‘eBayes’ and ‘topTable’ functions). FDRs were computed with empirical null distribution from the data using the t statistics value using fdrtool R package version 1.2.16. T-statistics were used as gene rank. Significant targets were defined as p value < 0.05. For individual gene plots we used variance stabilizing normalization (VSN) method from DEP R package version 1.6.1 to estimate protein abundance.

### TRAP and RNA-seq

TRAP-Seq was performed on hippocampi isolated from 4 littermate *Fmr1^-/y^* and wildtype mice hemizygous for *5nap25*-EGFPL10a, as previously described ^21^. For DHPG experiments, hippocampal slices were prepared from 10 CA1-TRAP *WT/Fmr1^-/y^* littermate pairs. For CX-5461 experiments, hippocampal slices were prepared from 9 CA1-TRAP *WT/Fmr1^-/y^* littermate pairs, with slices incubated with vehicle or 200nM CX-5461 for 30 min, stimulated with DHPG for 5 min (+/-CX-5461), and moved to fresh ACSF (+/-CX-5461) for an additional 25 min. For DHPG experiments, slices from 2 mice were pooled to obtain paired vehicle and DHPG samples from the same animals. For CX+DHPG experiments, slices from 3 mice were pooled to obtain paired vehicle, CX, DHPG and CX+DHPG conditions from the same animals. Samples were homogenized in ice-cold lysis buffer (20 mM HEPES, 5 mM MgCl_2_, 150 mM KCl, 0.5 mM DTT, 100 μg/ml cycloheximide, RNase inhibitors and protease inhibitors), and centrifuged at 1,000 x g for 10 minutes. Supernatants were then extracted with 1% NP-40 and 1% DHPC on ice and centrifuged at 20,000 x g for 20 minutes. The supernatant was incubated with streptavidin/protein L-coated Dynabeads bound to anti-GFP antibodies (HtzGFP-19F7 and HtzGFP-19C8, Memorial Sloan Kettering Centre) overnight at 4°C with gentle mixing to isolate translating ribosome-bound mRNA. Anti-GFP beads were washed with high salt buffer (20 mM HEPES, 5 mM MgCl_2_, 350 mM KCl, 1% NP-40, 0.5 mM DTT and 100 μg/ml cycloheximide) and RNA was eluted from all samples using PicoPure RNA isolation kit (Thermo-Fisher Scientific) according to the manufacturer’s instructions. RNA with RIN > 7 was prepared for sequencing using the RNAseq Ovation V2 kit (Nugen) or Smart-seq 4 low-abundance RNA kit (Takara) and sequenced on the Illumina HiSeq 4000 or Novaseq 2 platform in collaboration with Oxford Genomics Centre.

For downstream analysis, 75 bp or 150 bp paired-end reads were mapped to the *Mus musculus* primary assembly (Ensembl release v88) using STAR 2.4.0i. Reads that were uniquely aligned to annotated genes were counted with featureCounts 1.4.6-p2. Differential expression analyses were performed with a paired design to account for matching vehicle and DHPG stimulated samples, with fold change shrinkage using the normal prior (https://genomebiology.biomedcentral.com/articles/10.1186/s13059-014-0550-8), using DESeq2 version 1.18.1 (Bioconductor). WT and *Fmr1^-/y^* were analyzed separately. CA1 TRAP-Seq and hippocampal RNA-Seq datasets for *Fmr1^-/y^* were obtained from the 6 *Fmr1^-/y^* and WT littermate pairs (P25-32) used in our previous study, with datasets available at GEO:GSE101823 ^21^. The count matrix was re-analyzed with DESeq2 using the log2 fold change shrinkage using the normal prior to cancel out the bias for large fold change observed in genes with low expression level. Significance was determined using the standard DESeq2 FDR cut-off of adjusted p < 0.1, with full results shown in Tables.

### Gene set analysis

For generating gene sets, we obtained GO terms associated with each gene using BioMart. Ribosomal protein information was obtained from the Ribosomal Protein Gene Database (http://ribosome.med.miyazaki-u.ac.jp/). To assess the differential expression of RPs, z tests were performed between RPs and randomly selected genes of the same number detected in each dataset. Gene Set Enrichment Analysis (GSEA) ^24^ was used to calculate an enrichment score (ES) for each gene set in the ranked list of genes. This score is determined by a running-sum statistic that increases as a gene within the set is identified and decreases the score when a gene is absent. Significance of the ES was determined by comparison to a null distribution of ES scores calculated from 1000 permutations of scrambled gene lists. For population comparisons (e.g., RPs and 500 up/down comparisons), Gene Set mean rank Analysis (GSA mean) was calculated in a similar fashion, except the mean rank of all genes in a gene set was used in place of ES. To test the distribution of RPs, random sets containing the same number of genes as the ribosomal proteins were generated and used as null distributions. Nominal p-values for each gene set were corrected for multiple comparisons using FDR and both nominal and adjusted p-values are reported in Tables. All GSEA and GSA were performed with Piano package version 1.18.1 (Bioconductor) ^84^. For TRAP-Seq and RNA-Seq, moderated log_2_ fold change values were used as gene ranks. For proteomics, Bayes moderated effect sizes were used as gene ranks. Network plots were generated with igraph package version 1.1.2 in R using the number of shared genes as weights between the two points. GO enrichment analysis was performed with DAVID (https://pubmed.ncbi.nlm.nih.gov/19131956/). Clustering analyses were performed with igraph using edge betweenness algorithm. GO terms that share 90% genes were removed for the clustering analysis.

For length dependent gene set analysis, gene rank was set based on CDS length. Two different versions were created where the top rank was set to the shortest gene or the longest gene. GSA was performed based on the ranks with all cellular component GO terms, with the restriction of minimum 20 genes and maximum 50 genes, testing the mean difference for each GO term against the null distribution. The top 7 GO terms shown were selected based on p value and enrichment score.

### Synaptoneurosome preparation

Hippocampal slices were prepared as above and then homogenized in ice-cold homogenization buffer (10 mM HEPES, 2 mM EDTA, 2 mM EGTA, 150 mM NaCl) in 2 ml Dounce homogenizers. Samples were taken for immunoblotting, and the remaining homogenates were filtered through 2 x 100 μm filters (Millipore), followed by a 5 μm filter (Millipore). Samples were then centrifuged at 10,000 x g for 10 minutes and the pellets (synaptoneurosome sample) re-suspended in lysis buffer (50 mM HEPES, 5 mM EDTA, 150 mM NaCl, 1% Triton X-100, 0.5% sodium deoxycholate, 0.1% SDS). Protein concentrations were determined using BioRad DC kit (BioRad).

### Immunoblotting

Samples were boiled in Laemmli sample buffer and resolved using SDS-PAGE before being transferred to nitrocellulose and stained for total protein using the Memcode Reversible staining kit (Pierce). Membranes were blocked with 5% BSA in TBS + 0.1% Tween-20 for 1h, then incubated in primary antibody overnight at 4°C (Rpl10a Abcam 174318, Rps25 Thermo scientific PA5-56865). Membranes were then incubated with HRP-conjugated secondary antibodies for 30 minutes (Cell Signaling), developed with Clarity ECL (BioRad), and exposed to film. Densitometry was performed on scanned blot films and quantified using ImageStudio Lite (Licor). Densitometry data was normalized to total protein, which was quantified using scanned images of total protein memcode staining and quantified using FIJI.

All samples were loaded blind to genotype and condition, and were thus randomized across the gel. This was done to prevent skewing that can occur when all samples of one condition are loaded towards the ends of the gel. For figures, different lanes from the same gel are shown next to one another with blank space to indicate they are not loaded side-by-side, and unprocessed blots are shown in the Supplementary Figures. To correct for blot-to-blot variance, each blot signal was normalized to the average signal of all lanes on the same blot. All gels were analyzed blind to genotype.

### Flow cytometry

Hippocampal slices were prepared and recovered as above. The CA1 was micro-dissected and incubated in ACSF with papain (20 U/ml; Sigma-Aldrich) for 45 minutes at 37°C with 5% CO_2_. The tissue was dissociated using a fire-polished glass pipette and fixed with 4 % PFA. The tissue dissociate was then filtered with a 70 μm cell sieve and blocked with 1.5% FCS in PBS with 0.1% saponin, prior to overnight incubation at 4°C with a Rpl10a antibody (Abcam, ab174318). Alexa Fluor 594 conjugated secondary antibody (Thermo Fisher) and Alexa Fluor 488 conjugated anti-NeuN (Abcam, ab190195) were applied for 1 hour at room temperature. Flow analyses were performed using the LSRFortessa (BD bioscience) and the data analyzed using FlowJo software in collaboration with the QMRI flow cytometry core facility at the University of Edinburgh. To correct for the experiment-to-experiment signal intensity variance, each value obtained in an experiment was normalized by the average value obtained from all cells in that experiment regardless of the genotype. All staining and analysis were performed blind to genotype.

### Immunostaining

Mice were perfused with 4% PFA and 50 μm coronal sections were collected from the dorsal hippocampus. Immunostaining was performed blind to genotype on littermate *Fmr1^-/y^* and WT pairs using Y10b (Abcam, ab171119) and NeuN (Merck, ABN78) or Fibrillarin (Abcam, ab5821) and NeuN (Millipore, MAB377) antibodies. All imaging was performed on a Zeiss LSM800 confocal microscope in collaboration with the IMPACT facility at the University of Edinburgh. The edges of single neurons were set manually across each z plane based on NeuN (for Y10b signal) or DAPI (for Fibrillarin) staining. Y10b or Fibrillarin volume within the defined area was then reconstructed with automated settings from IMARIS 9.1.7/9.2.0. Cytoplasmic rRNA expression was calculated as the Y10b volume per NeuN volume. To control for variation in immunostaining, yoked pairs of *Fmr1^-/y^* and WT littermates were processed together, and calculated volumes were normalized to the average volume of all neurons regardless of genotype for each experimental session.

### RT-qPCR

RNA for each sample was converted into cDNA using Superscript VILO cDNA Synthesis Kit (Life Technologies) and RT-qPCR was performed using Quantitect SYBRgreen qPCR master mix (Qiagen) according to the manufacturer’s instructions. Samples were prepared in triplicate in 96-well reaction plates and run on a StepOne Plus (Life Technologies). Primers used for RT-qPCR are as follows: *b-actin* (F-CACCACACCTTCTACAATGAG, R- GTCTCAAACATGATCTGGGTC), *Ppib1* (F- CAGCAAGTTCCATCGTGTCA, R- GATGCTCTTTCCTCCTGTGC), *Npas4* (F- CAGGGACAGGTTAGGGTTCA, R- TTCAGCAGATCAGCCAGTTG), *Arc* (F- CAGGGGTGAGCTGAAGCCACAAA, R- CCATGTAGGCAGCTTCAGGAGAAGAGAG), *Rps25* (F- GCTCTGTAAGGAGGTTCCGA, R- CGTCCCCACCCTTTGTGTTT), *Camk2a* (F-GGAATCTTCTGAGAGCACCA, R-CACATCTTCGTGTAGGACTC). Fold change was calculated using the delta-delta Ct method where the first delta was calculated as the average of *b-actin* and *Ppib1*, and the second delta was calculated as the average of all samples run together for each experiment to avoid cancelling out the variation in control samples. Values were then normalized to the mean of all control samples for graphical purposes.

### Transcript length analysis

Total transcript length, CDS length and the CDS sequences were obtained from BioMart. For each gene, the most abundant transcript in the hippocampus was determined by assessing the transcript expression level from total hippocampal RNA-Seq (GEO:GSE101823) using TPM values. Information for the most abundant transcript was used for all downstream analysis. GC content of the CDS sequence was calculated using seqinr package version 3.4-5 (Bioconductor). For *Fmr1^-/y^* length analysis, in order to compare this to the available RNA-Seq dataset from the hippocampus we restricted the analysis to previously determined high confidence (>2 FPKM) dorsal CA1 pyr neuron expressing genes ^85^. For total RNA abundance correction, DESeq2 normalized counts were obtained from our previous work ^21^ and we took the fraction of TRAP-Seq against the matched RNA-Seq sample. For top regulated genes, we took the genes that were up or downregulated in all 6 pairs.

### 5’UTR sequence

To obtain an accurate 5’UTR sequence, the transcription start site (TSS) was determined by analyzing a published CAGE-Seq dataset from FANTOM database (FF:13-16E8) using a custom Python script. Briefly, each annotated protein coding gene was scanned from 1000 bases upstream of the earliest start location of any transcript to find the greatest pile up of reads and this was determined as the transcription start site. Then the transcript was chosen for which this site was the smallest number of bases upstream of the coding start location, and the 5’ UTR sequence defined as the region between the transcription start site and the transcript coding start location. 5’ UTRs longer than 500 bases, and genes for which the defined transcription start site was downstream of any coding start location, were excluded.

### SFARI genes

SFARI genes were identified from the SFARI GENE database (https://gene.sfari.org/xg, up to date on September 2018), and those with a gene score of 1-4 used for downstream analysis. High-confidence SFARI genes were defined by a gene score of 1-2. A z-test was used to test the difference in the mean of the distributions. For CDS length vs gene length analysis, threshold for CDS length group was set at log2(CDS length) = 11 and the threshold for gene length was set at log2(gene length) = 16.

### FMRP target analysis

The stringent FMRP target list was obtained from a previously published study looking at cell type specific FMRP targets in CA1 pyramidal neurons ^58^. A two-sample z-test was used to test the difference in the mean of the distributions between FMRP targets against all detected transcripts within the population.

### LTD functional term analysis

GO terms that are down regulated with adjusted p < 0.01 in WT DHPG experiments were denoted LTD down GO terms. For each of the LTD down GO terms, the z score in the WT CX+DHPG condition or the *Fmr1^-/y^* CX+DHPG condition was calculated by the formula below.

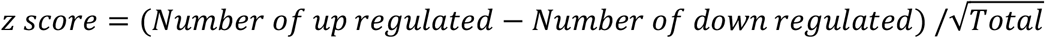

The distribution of LTD down GO terms was compared to a null distribution of z-scores of all GO terms detected in the comparison with a z-test.

### Electrophysiology

Horizontal hippocampal slices (400 μM) were prepared from *Fmr1^-/y^* and WT littermates (P25-32), blind to genotype, in ice-cold dissection buffer (86 mM NaCl, 25 mM KCl, 1.2 mM NaH_2_PO_4_, 25 mM NaHCO_3_, mM 20 glucose, 0.5 mM CaCl_2_, 7 mM MgCl_2_, saturated with 95 % O_2_ and 5 % CO_2_) and an incision made through CA3. Slices were recovered for at least 2 hours at 30 °C in ACSF (124 mM NaCl, 5 mM KCl, 1.25 mM NaH_2_PO_4_, 26 mM NaHCO_3_, 10 mM glucose, 2 mM CaCl_2_; 1 mM MgCl_2_, saturated with 95 % O_2_ and 5 % CO_2_) before being transferred to a submersion chamber heated to 30 °C and perfused with ACSF containing either DMSO vehicle or CX-5461 (200 nM) for at least 30 mins before recording. Field excitatory postsynaptic potentials (fEPSPs) were evoked by applying a current pulse to the Schaffer collateral pathway every 30 s with a bipolar stimulating electrode and recording with an extracellular electrode (1-3 MΩ) in stratum radiatum of hippocampal CA1. Following a 20-minute stable baseline, LTD was induced by the application of S-DHPG (50 μM; 5 min) in the presence of either vehicle (0.002% DMSO in ddH_2_O) or CX-5461 (200 nM), which was present for the duration of the recording (55 minutes post DHPG washout). The magnitude of LTD was calculated from average fEPSP slope during the last 10 minutes of recording relative to fEPSP slope during the 20-minute baseline. All recordings were completed within 7 hours of slice preparation. Data were analyzed blind to genotype and treatment, with unstable recordings (baseline drift +/- 5%) filtered out.

### Quantification and statistical analysis

All statistics were performed using R or Graphpad Prism v9. For RNA-seq datasets, differential expression was determined using DESeq2 using the default cutoff for significance (adjusted p-value < 0.1) unless otherwise noted. For GSEA, significance was determined by nominal p-value based on comparisons to a permutation-generated null distribution, and adjusted p-values were generated to correct for multiple comparisons with FDR cut off as indicated. For immunoblotting, immunostaining and qPCR experiments, significance between more than 2 groups was determined by two-way mixed-model ANOVA followed by post-hoc tests (Sidak’s or FDR as indicated). For comparison of two groups of yoked *Fmr1^-/y^* and WT samples, prepared and analyzed in the same experiment (i.e., sliced in the same experiment and run on the same gel or imaged in the same session), significance was determined by paired t-test. For correlation analysis, significance was determined by Pearson’s correlation coefficient. For analysis of transcript characteristics Wilcoxon rank sum tests were used to compare up/down groups to all detected genes. Differences between distributions were compared using KS test or two-sample z-test as indicated. Overlap analyses were performed using hypergeometric test.

## Data Availability

All sequencing and proteomics data will be deposited in GEO (https://www.ncbi.nlm.nih.gov/geo/) or similar repository upon publication. All renewable reagents and protocols will be available upon reasonable request.

## Code Availability

All custom scripts will be deposited upon publication.

## Acknowledgments

Special thanks to David JA Wyllie for help overseeing LTD recordings, and to Katherine Bonnycastle and Mike A Cousin for providing excellent advice for the project. Thanks to Serena Linley-Adams, Katy Homyer, Keiron Scrimger and Hannah Wat for help with image analysis. Confocal imaging and IMARIS reconstruction were performed in collaboration with the IMPACT facility at the University of Edinburgh.

The authors are grateful for support from the Wellcome Trust/Royal Society (SHDF 104116/Z/14/Z and SRF 219556/Z/19/Z), Medical Research Council (MRC MR/M006336/1 and MR/S026312/1), and Simons Initiative for the Developing Brain (SIDB).

## Author contributions

EKO and SSS conceptualized the study and prepared the manuscript. SRL prepared tissue for proteomics and performed all biochemistry experiments. SSS, NCV and SRT performed TRAP experiments. SSS performed all bioinformatics analyses with assistance from OD. SSS and NA performed mGluR-LTD recordings with supervision of EKO and PCK. MAGL performed Mass Spectroscopy experiments with supervision of KWL, and analyzed results with SSS. SSS and CBH performed confocal imaging analyses. JCD provided essential insights and advice on the study and manuscript.

## Supplementary Information

**Supplementary Figures 1-5 contain information related to Figures 1-6**

**Supplementary Tables 1-13 contain raw data from proteomics and RNA-seq, and results from GSEA and overlap analyses**

## References

1 Kelleher, R. J., 3rd & Bear, M. F. The autistic neuron: troubled translation? Cell 135, 401–406, doi:10.1016/j.cell.2008.10.017 (2008).

2 Louros, S. R. & Osterweil, E. K. Perturbed proteostasis in autism spectrum disorders. J Neurochem, doi:10.1111/jnc.13723 (2016).

3 Darnell, J. C. et al. FMRP stalls ribosomal translocation on mRNAs linked to synaptic function and autism. Cell 146, 247–261, doi:S0092-8674(11)00655-6[pii]10.1016/j.cell.2011.06.013 [doi] (2011).

4 Osterweil, E. K., Krueger, D. D., Reinhold, K. & Bear, M. F. Hypersensitivity to mGluR5 and ERK1/2 leads to excessive protein synthesis in the hippocampus of a mouse model of fragile X syndrome. J Neurosci 30, 15616–15627 (2010).

5 Qin, M., Kang, J., Burlin, T. V., Jiang, C. & Smith, C. B. Postadolescent changes in regional cerebral protein synthesis: an in vivo study in the FMR1 null mouse. J Neurosci 25, 5087–5095 (2005).

6 Bolduc, F. V., Bell, K., Cox, H., Broadie, K. S. & Tully, T. Excess protein synthesis in Drosophila fragile X mutants impairs long-term memory. Nat Neurosci 11, 1143–1145 (2008).

7 Till, S. M. et al. Conserved hippocampal cellular pathophysiology but distinct behavioural deficits in a new rat model of FXS. Human molecular genetics 24, 5977–5984, doi:10.1093/hmg/ddv299 (2015).

8 Stoppel, L. J., Osterweil, E. K. & Bear, M. F. in Fragile X Syndrome: From Genetics to Targeted Treatment (eds R. Willemsen & F. Kooy) Ch. 9, (Elsevier, 2017).

9 Richter, J. D., Bassell, G. J. & Klann, E. Dysregulation and restoration of translational homeostasis in fragile X syndrome. Nat Rev Neurosci 16, 595–605, doi:10.1038/nrn4001 (2015).

10 Berry-Kravis, E. M. et al. Drug development for neurodevelopmental disorders: lessons learned from fragile X syndrome. Nature reviews. Drug discovery 17, 280–299, doi:10.1038/nrd.2017.221 (2018).

11 Das Sharma, S. et al. Widespread Alterations in Translation Elongation in the Brain of Juvenile Fmr1 Knockout Mice. Cell Rep 26, 3313–3322 e3315, doi:10.1016/j.celrep.2019.02.086 (2019).

12 Shah, S. et al. FMRP Control of Ribosome Translocation Promotes Chromatin Modifications and Alternative Splicing of Neuronal Genes Linked to Autism. Cell Rep 30, 4459–4472 e4456, doi: 10.1016/j.celrep.2020.02.076 (2020).

13 Huber, K. M., Gallagher, S. M., Warren, S. T. & Bear, M. F. Altered synaptic plasticity in a mouse model of fragile X mental retardation. Proc Natl Acad Sci U S A 99, 7746–7750 (2002).

14 Bear, M. F., Huber, K. M. & Warren, S. T. The mGluR theory of fragile X mental retardation. Trends in neurosciences 27, 370–377, doi:10.1016/j.tins.2004.04.009 [doi] S0166-2236(04)00132-8 [pii] (2004).

15 Bhakar, A. L., Dolen, G. & Bear, M. F. The pathophysiology of fragile X (and what it teaches us about synapses). Annual review of neuroscience 35, 417–443, doi:10.1146/annurev-neuro-060909-153138 [doi] (2012).

16 Huber, K. M., Kayser, M. S. & Bear, M. F. Role for rapid dendritic protein synthesis in hippocampal mGluR-dependent long-term depression. Science 288, 1254–1257 (2000).

17 Nosyreva, E. D. & Huber, K. M. Metabotropic receptor-dependent long-term depression persists in the absence of protein synthesis in the mouse model of fragile X syndrome. J Neurophysiol 95, 3291–3295 (2006).

18 Dolen, G. et al. Correction of fragile X syndrome in mice. Neuron 56, 955–962, doi: 10.1016/j.neuron.2007.12.001 (2007).

19 Zalfa, F. et al. Fragile X mental retardation protein (FMRP) binds specifically to the brain cytoplasmic RNAs BC1/BC200 via a novel RNA-binding motif. J Biol Chem 280, 33403–33410 (2005).

20 Jakkamsetti, V. et al. Experience-induced Arc/Arg3.1 primes CA1 pyramidal neurons for metabotropic glutamate receptor-dependent long-term synaptic depression. Neuron 80, 72–79, doi:S0896-6273(13)00644-2 [pii] 10.1016/j.neuron.2013.07.020 [doi] (2013).

21 Thomson, S. R. et al. Cell-Type-Specific Translation Profiling Reveals a Novel Strategy for Treating Fragile X Syndrome. Neuron 95, 550–563 e555, doi:10.1016/j.neuron.2017.07.013 (2017).

22 Tang, B. et al. Fmr1 deficiency promotes age-dependent alterations in the cortical synaptic proteome. Proc Natl Acad Sci U S A 112, E4697–4706, doi: 10.1073/pnas.1502258112 (2015).

23 Stasyk, T. & Huber, L. A. Zooming in: fractionation strategies in proteomics. Proteomics 4, 3704–3716, doi: 10.1002/pmic.200401048 (2004).

24 Subramanian, A. et al. Gene set enrichment analysis: a knowledge-based approach for interpreting genome-wide expression profiles. Proc Natl Acad Sci U S A 102, 15545–15550, doi:10.1073/pnas.0506580102 (2005).

25 Lafontaine, D. L. & Tollervey, D. The function and synthesis of ribosomes. Nature reviews. Molecular cell biology 2, 514–520, doi: 10.1038/35080045 (2001).

26 Shi, Z. et al. Heterogeneous Ribosomes Preferentially Translate Distinct Subpools of mRNAs Genome-wide. Mol Cell 67, 71–83 e77, doi:10.1016/j.molcel.2017.05.021 (2017).

27 Sharifi, S. & Bierhoff, H. Regulation of RNA Polymerase I Transcription in Development, Disease, and Aging. Annu Rev Biochem 87, 51–73, doi: 10.1146/annurev-biochem-062917-012612 (2018).

28 Kim, H. K., Kim, Y. B., Kim, E. G. & Schuman, E. Measurement of dendritic mRNA transport using ribosomal markers. Biochem Biophys Res Commun 328, 895–900, doi:10.1016/j.bbrc.2005.01.041 (2005).

29 Muddashetty, R. S., Kelic, S., Gross, C., Xu, M. & Bassell, G. J. Dysregulated metabotropic glutamate receptor-dependent translation of AMPA receptor and postsynaptic density-95 mRNAs at synapses in a mouse model of fragile X syndrome. J Neurosci 27, 5338–5348 (2007).

30 Di Prisco, G. V. et al. Translational control of mGluR-dependent long-term depression and object-place learning by eIF2alpha. Nat Neurosci 17, 1073–1082, doi: 10.1038/nn.3754 (2014).

31 Bowling, H. et al. Altered steady state and activity-dependent de novo protein expression in fragile X syndrome. Nat Commun 10, 1710, doi:10.1038/s41467-019-09553-8 (2019).

32 Huber, K. M., Roder, J. C. & Bear, M. F. Chemical induction of mGluR5- and protein synthesis--dependent long-term depression in hippocampal area CA1. J Neurophysiol 86, 321–325 (2001).

33 Sun, X. & Lin, Y. Npas4: Linking Neuronal Activity to Memory. Trends in neurosciences 39, 264–275, doi:10.1016/j.tins.2016.02.003 (2016).

34 Cole, A. J., Saffen, D. W., Baraban, J. M. & Worley, P. F. Rapid increase of an immediate early gene messenger RNA in hippocampal neurons by synaptic NMDA receptor activation. Nature 340, 474–476, doi:10.1038/340474a0 (1989).

35 Fusco, C. M. et al. Neuronal ribosomes exhibit dynamic and context-dependent exchange of ribosomal proteins. Nat Commun 12, 6127, doi:10.1038/s41467-021-26365-x (2021).

36 Andreassi, C. et al. An NGF-responsive element targets myo-inositol monophosphatase-1 mRNA to sympathetic neuron axons. Nat Neurosci 13, 291–301, doi: 10.1038/nn.2486 (2010).

37 Cajigas, I. J. et al. The local transcriptome in the synaptic neuropil revealed by deep sequencing and high-resolution imaging. Neuron 74, 453–466, doi:10.1016/j.neuron.2012.02.036 (2012).

38 Saal, L., Briese, M., Kneitz, S., Glinka, M. & Sendtner, M. Subcellular transcriptome alterations in a cell culture model of spinal muscular atrophy point to widespread defects in axonal growth and presynaptic differentiation. Rna 20, 1789–1802, doi: 10.1261/rna.047373.114 (2014).

39 Shigeoka, T. et al. On-Site Ribosome Remodeling by Locally Synthesized Ribosomal Proteins in Axons. Cell Rep 29, 3605–3619 e3610, doi:10.1016/j.celrep.2019.11.025 (2019).

40 Petibon, C., Malik Ghulam, M., Catala, M. & Abou Elela, S. Regulation of ribosomal protein genes: An ordered anarchy. Wiley Interdiscip Rev RNA 12, e1632, doi: 10.1002/wrna.1632 (2021).

41 Drygin, D. et al. Targeting RNA polymerase I with an oral small molecule CX-5461 inhibits ribosomal RNA synthesis and solid tumor growth. Cancer Res 71, 1418–1430, doi:10.1158/0008-5472.CAN-10-1728 (2011).

42 Khajuria, R. K. et al. Ribosome Levels Selectively Regulate Translation and Lineage Commitment in Human Hematopoiesis. Cell 173, 90–103 e119, doi:10.1016/j.cell.2018.02.036 (2018).

43 Allen, K. D. et al. Nucleolar integrity is required for the maintenance of long-term synaptic plasticity. PLoS One 9, e104364, doi:10.1371/journal.pone.0104364 (2014).

44 Zylka, M. J., Simon, J. M. & Philpot, B. D. Gene length matters in neurons. Neuron 86, 353–355, doi:10.1016/j.neuron.2015.03.059 (2015).

45 Eisenberg, E. & Levanon, E. Y. Human housekeeping genes are compact. Trends Genet 19, 362–365, doi:10.1016/S0168-9525(03)00140-9 (2003).

46 Arava, Y. et al. Genome-wide analysis of mRNA translation profiles in Saccharomyces cerevisiae. Proc Natl Acad Sci U S A 100, 3889–3894, doi: 10.1073/pnas.0635171100 (2003).

47 Fernandes, L. D., Moura, A. P. S. & Ciandrini, L. Gene length as a regulator for ribosome recruitment and protein synthesis: theoretical insights. Sci Rep 7, 17409, doi:10.1038/s41598-017-17618-1 (2017).

48 Ingolia, N. T., Ghaemmaghami, S., Newman, J. R. & Weissman, J. S. Genome-wide analysis in vivo of translation with nucleotide resolution using ribosome profiling. Science 324, 218–223, doi: 10.1126/science.1168978 (2009).

49 Lodish, H. F. Model for the regulation of mRNA translation applied to haemoglobin synthesis. Nature 251, 385–388, doi:10.1038/251385a0 (1974).

50 Shah, P., Ding, Y., Niemczyk, M., Kudla, G. & Plotkin, J. B. Rate-limiting steps in yeast protein translation. Cell 153, 1589–1601, doi:10.1016/j.cell.2013.05.049 (2013).

51 Mills, E. W. & Green, R. Ribosomopathies: There’s strength in numbers. Science 358, doi: 10.1126/science.aan2755 (2017).

52 Gorochowski, T. E., Avcilar-Kucukgoze, I., Bovenberg, R. A., Roubos, J. A. & Ignatova, Z. A Minimal Model of Ribosome Allocation Dynamics Captures Trade-offs in Expression between Endogenous and Synthetic Genes. ACS Synth Biol 5, 710–720, doi:10.1021/acssynbio.6b00040 (2016).

53 Oshlack, A. & Wakefield, M. J. Transcript length bias in RNA-seq data confounds systems biology. Biol Direct 4, 14, doi:10.1186/1745-6150-4-14 (2009).

54 King, I. F. et al. Topoisomerases facilitate transcription of long genes linked to autism. Nature 501, 58–62, doi:10.1038/nature12504 (2013).

55 Shohat, S. & Shifman, S. Bias towards large genes in autism. Nature 512, E1–2, doi:10.1038/nature13583 (2014).

56 Duda, M. et al. Brain-specific functional relationship networks inform autism spectrum disorder gene prediction. Transl Psychiatry 8, 56, doi: 10.1038/s41398-018-0098-6 (2018).

57 Ceolin, L. et al. Cell Type-Specific mRNA Dysregulation in Hippocampal CA1 Pyramidal Neurons of the Fragile X Syndrome Mouse Model. Front Mol Neurosci 10, 340, doi:10.3389/fnmol.2017.00340 (2017).

58 Sawicka, K. et al. FMRP has a cell-type-specific role in CA1 pyramidal neurons to regulate autism-related transcripts and circadian memory. eLife 8, doi:10.7554/eLife.46919 (2019).

59 Aryal, S., Longo, F. & Klann, E. Genetic removal of p70 S6K1 corrects coding sequence length-dependent alterations in mRNA translation in fragile X syndrome mice. Proc Natl Acad Sci U S A 118, doi: 10.1073/pnas.2001681118 (2021).

60 Niswender, C. M. & Conn, P. J. Metabotropic glutamate receptors: physiology, pharmacology, and disease. Annu Rev Pharmacol Toxicol 50, 295–322, doi: 10.1146/annurev.pharmtox.011008.145533 (2010).

61 Osterweil, E. K. et al. Lovastatin corrects excess protein synthesis and prevents epileptogenesis in a mouse model of fragile X syndrome. Neuron 77, 243–250 (2013).

62 Michalon, A. et al. Chronic Pharmacological mGlu5 Inhibition Corrects Fragile X in Adult Mice. Neuron 74, 49–56 (2012).

63 Chen, E., Sharma, M. R., Shi, X., Agrawal, R. K. & Joseph, S. Fragile X mental retardation protein regulates translation by binding directly to the ribosome. Mol Cell 54, 407–417, doi:10.1016/j.molcel.2014.03.023 (2014).

64 Banko, J. L., Hou, L., Poulin, F., Sonenberg, N. & Klann, E. Regulation of eukaryotic initiation factor 4E by converging signaling pathways during metabotropic glutamate receptor-dependent long-term depression. J Neurosci 26, 2167–2173 (2006).

65 Mathis, A. D. et al. Mechanisms of In Vivo Ribosome Maintenance Change in Response to Nutrient Signals. Mol Cell Proteomics 16, 243–254, doi: 10.1074/mcp.M116.063255 (2017).

66 Dorrbaum, A. R., Alvarez-Castelao, B., Nassim-Assir, B., Langer, J. D. & Schuman, E. M. Proteome dynamics during homeostatic scaling in cultured neurons. eLife 9, doi: 10.7554/eLife.52939 (2020).

67 Brackmann, M., Zhao, C., Kuhl, D., Manahan-Vaughan, D. & Braunewell, K. H. MGluRs regulate the expression of neuronal calcium sensor proteins NCS-1 and VILIP-1 and the immediate early gene arg3.1/arc in the hippocampus in vivo. Biochem Biophys Res Commun 322, 1073–1079, doi:10.1016/j.bbrc.2004.08.028 (2004).

68 Shepherd, J. D. et al. Arc/Arg3.1 mediates homeostatic synaptic scaling of AMPA receptors. Neuron 52, 475–484 (2006).

69 Wang, H. & Zhuo, M. Group I metabotropic glutamate receptor-mediated gene transcription and implications for synaptic plasticity and diseases. Front Pharmacol 3, 189, doi:10.3389/fphar.2012.00189 (2012).

70 Liu, F. et al. Regulation of cyclin-dependent kinase 5 and casein kinase 1 by metabotropic glutamate receptors. Proc Natl Acad Sci U S A 98, 11062–11068, doi: 10.1073/pnas.191353898 (2001).

71 Comery, T. A. et al. Abnormal dendritic spines in fragile X knockout mice: maturation and pruning deficits. Proc Natl Acad Sci U S A 94, 5401–5404 (1997).

72 Vanderklish, P. W. & Edelman, G. M. Dendritic spines elongate after stimulation of group 1 metabotropic glutamate receptors in cultured hippocampal neurons. Proc Natl Acad Sci U S A 99, 1639–1644, doi:10.1073/pnas.032681099 [doi] 032681099 [pii] (2002).

73 He, C. X. & Portera-Cailliau, C. The trouble with spines in fragile X syndrome: density, maturity and plasticity. Neuroscience 251, 120–128, doi:10.1016/j.neuroscience.2012.03.049 (2013).

74 Pasterkamp, R. J. Getting neural circuits into shape with semaphorins. Nat Rev Neurosci 13, 605–618, doi:10.1038/nrn3302 (2012).

75 Shu, H. et al. FMRP links optimal codons to mRNA stability in neurons. Proc Natl Acad Sci U S A 117, 30400–30411, doi:10.1073/pnas.2009161117 (2020).

76 Greenblatt, E. J. & Spradling, A. C. Fragile X mental retardation 1 gene enhances the translation of large autism-related proteins. Science 361, 709–712, doi: 10.1126/science.aas9963 (2018).

77 Accogli, A. et al. De Novo Pathogenic Variants in N-cadherin Cause a Syndromic Neurodevelopmental Disorder with Corpus Collosum, Axon, Cardiac, Ocular, and Genital Defects. American journal of human genetics 105, 854–868, doi:10.1016/j.ajhg.2019.09.005 (2019).

78 Braat, S. & Kooy, R. F. The GABAA Receptor as a Therapeutic Target for Neurodevelopmental Disorders. Neuron 86, 1119–1130, doi:10.1016/j.neuron.2015.03.042 (2015).

79 Contractor, A., Klyachko, V. A. & Portera-Cailliau, C. Altered Neuronal and Circuit Excitability in Fragile X Syndrome. Neuron 87, 699–715, doi:10.1016/j.neuron.2015.06.017 (2015).

80 Maurin, T. et al. Involvement of Phosphodiesterase 2A Activity in the Pathophysiology of Fragile X Syndrome. Cerebral cortex 29, 3241–3252, doi: 10.1093/cercor/bhy192 (2019).

81 Deng, P. Y. et al. FMRP regulates neurotransmitter release and synaptic information transmission by modulating action potential duration via BK channels. Neuron 77, 696–711, doi:10.1016/j.neuron.2012.12.018 (2013).

82 Koopmans, F. et al. Comparative Hippocampal Synaptic Proteomes of Rodents and Primates: Differences in Neuroplasticity-Related Proteins. Front Mol Neurosci 11, 364, doi: 10.3389/fnmol.2018.00364 (2018).

83 Koopmans, F., Ho, J. T. C., Smit, A. B. & Li, K. W. Comparative Analyses of Data Independent Acquisition Mass Spectrometric Approaches: DIA, WiSIM-DIA, and Untargeted DIA. Proteomics 18, doi:10.1002/pmic.201700304 (2018).

84 Varemo, L., Nielsen, J. & Nookaew, I. Enriching the gene set analysis of genome-wide data by incorporating directionality of gene expression and combining statistical hypotheses and methods. Nucleic Acids Res 41, 4378–4391, doi: 10.1093/nar/gkt111 (2013).

85 Cembrowski, M. S., Wang, L., Sugino, K., Shields, B. C. & Spruston, N. Hipposeq: a comprehensive RNA-seq database of gene expression in hippocampal principal neurons. eLife 5, e14997, doi:10.7554/eLife.14997 (2016).

