## Supplemental Information for "Excess ribosomal protein production unbalances translation in Fragile X Syndrome"

* Equal contributions

**Supplementary Information:**

**Supplementary Table 1.** Contains raw data from the *Fmr1*$^{-/y}$ proteomics analysis
**Supplementary Table 2.** Contains GSEA analyses of TRAP-seq and proteomics datasets
**Supplementary Table 3.** Contains raw data from *Fmr1*$^{-/y}$ SNAP-TRAP-seq
**Supplementary Table 4.** Contains raw data from DHPG TRAP-seq
**Supplementary Table 5.** Contains GSEA analyses of DHPG TRAP-seq
**Supplementary Table 6.** Lists targets upregulated in both WT DHPG TRAP-seq and in *Fmr1*$^{-/y}$ CA1-TRAP-seq
**Supplementary Table 7.** Lists RPs upregulated in both WT DHPG TRAP-seq and in *Fmr1*$^{-/y}$ CA1-TRAP-seq
**Supplementary Table 8.** Contains GSEA analyses of transcript lengths in TRAP-seq
**Supplementary Table 9.** Lists targets downregulated in both *Fmr1*$^{-/y}$ proteomics and *Fmr1*$^{-/y}$ SNAP-TRAP-seq datasets
**Supplementary Table 10.** Lists targets downregulated in both *Fmr1*$^{-/y}$ proteomics and *Fmr1*$^{-/y}$ CA1-TRAP-seq datasets
**Supplementary Table 11.** Contains raw data from DHPG transcriptome
**Supplementary Table 12.** Lists targets downregulated in both WT DHPG TRAP-seq and in *Fmr1*$^{-/y}$ CA1-TRAP-seq
**Supplementary Table 13.** Contains raw data from CX-5461 DHPG TRAP-seq

**Supplementary Figures 1-5**

**Supplementary Figure 1**

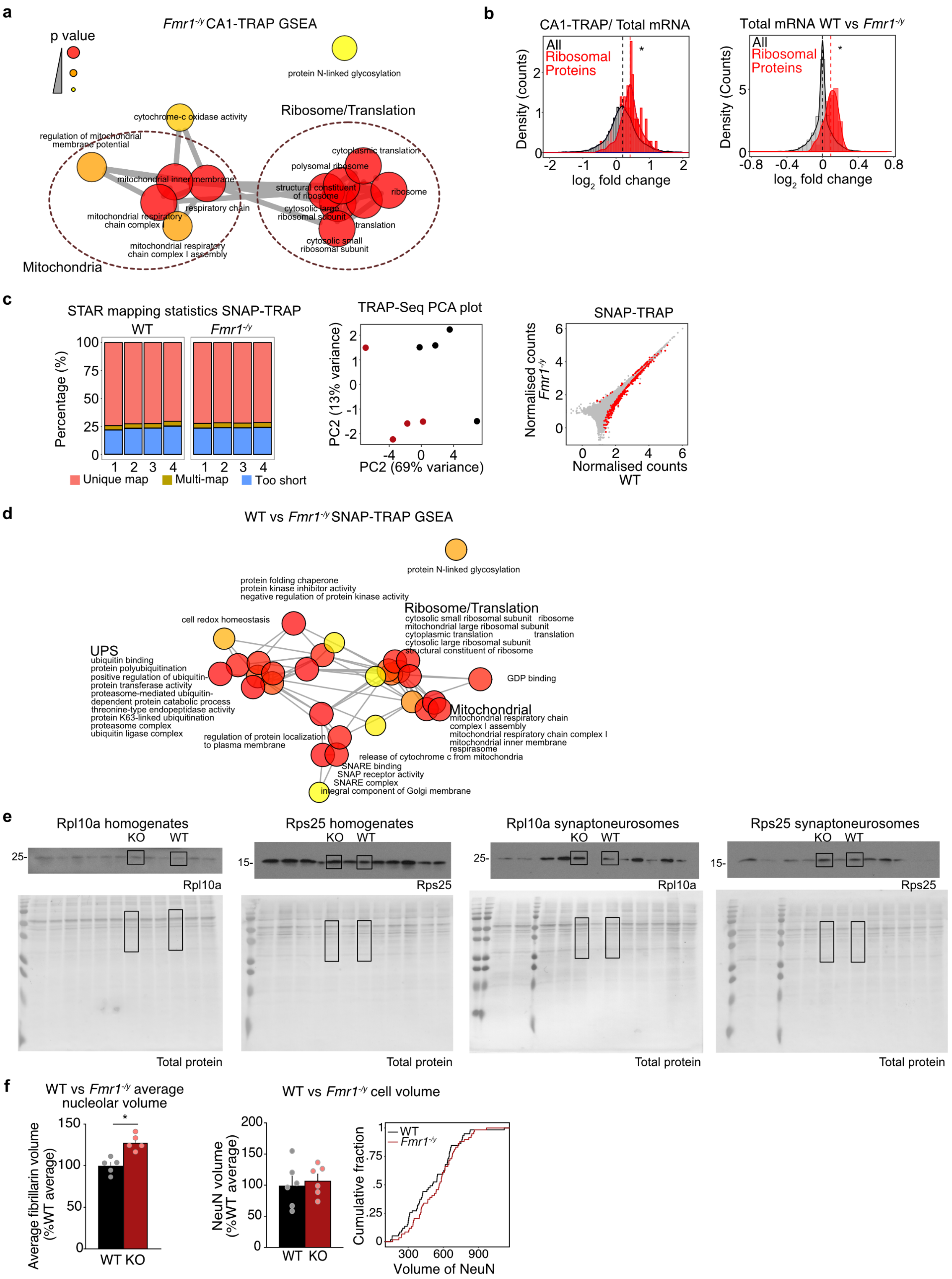


**Supplementary Figure 1. Ribosome upregulation in *Fmr1*$^{-/y}$ neurons. (a-b)** Similar to what is seen in the *Fmr1*$^{-/y}$ proteome, network analysis of upregulated GO terms in *Fmr1*$^{-/y}$ CA1 TRAP-Seq (adjusted p value $< 0.1$) and SNAP TRAP-Seq (adjusted p value $< 0.01$) identifies ribosome and translation related GO terms as a prominent cluster. Node size and color denotes significance and thickness of lines denotes the number of genes shared between nodes. **(c)** The increased RP expression persists after normalizing the CA1-TRAP-Seq to RNA-seq population (two sample z test, $z = 2.16$, $p = 0.031$), suggesting an increase in translation. However, a significant increase in RP transcripts is also seen in the total mRNA population of *Fmr1*$^{-/y}$ hippocampus (two sample z test, z = 7.47, p =7.78X10$^{-14}$), indicating that abundance is also increased. (**d**) Mapping statistics and PCA plot for the SNAP-TRAP samples are shown, as are the DESeq2 results with significant genes detected in red. (**e**) Raw immunoblots and total protein memcode staining used to quantify Rpl10a and Rps25 expression in **Fig. 1** are shown. As samples are loaded blind to genotype, the arrangement is random and must be re-ordered for the main figure with spaces denoting lanes that are not run next to each other. (**f**) Similar to results seen with total nucleolar volume in *Fmr1*$^{-/y}$ neurons, calculation of the average nucleolar volume per neuron reveals a significant increase versus WT (paired t test, WT = 100 ± 4.235%, *Fmr1*$^{-/y}$ = 128 ± 4.235%, * p = 0.0296, N = 5 littermate pairs). No significant difference was observed in reconstructed NeuN volume of *Fmr1*$^{-/y}$ versus WT neurons (paired t test, WT = 100 ± 14.49%, *Fmr1*$^{-/y}$ = 107.9 ± 10.0%, p = 0.6072, N= 6 animals, KS test p = 0.65, N= 59 cells).

**Supplementary Figure 2**

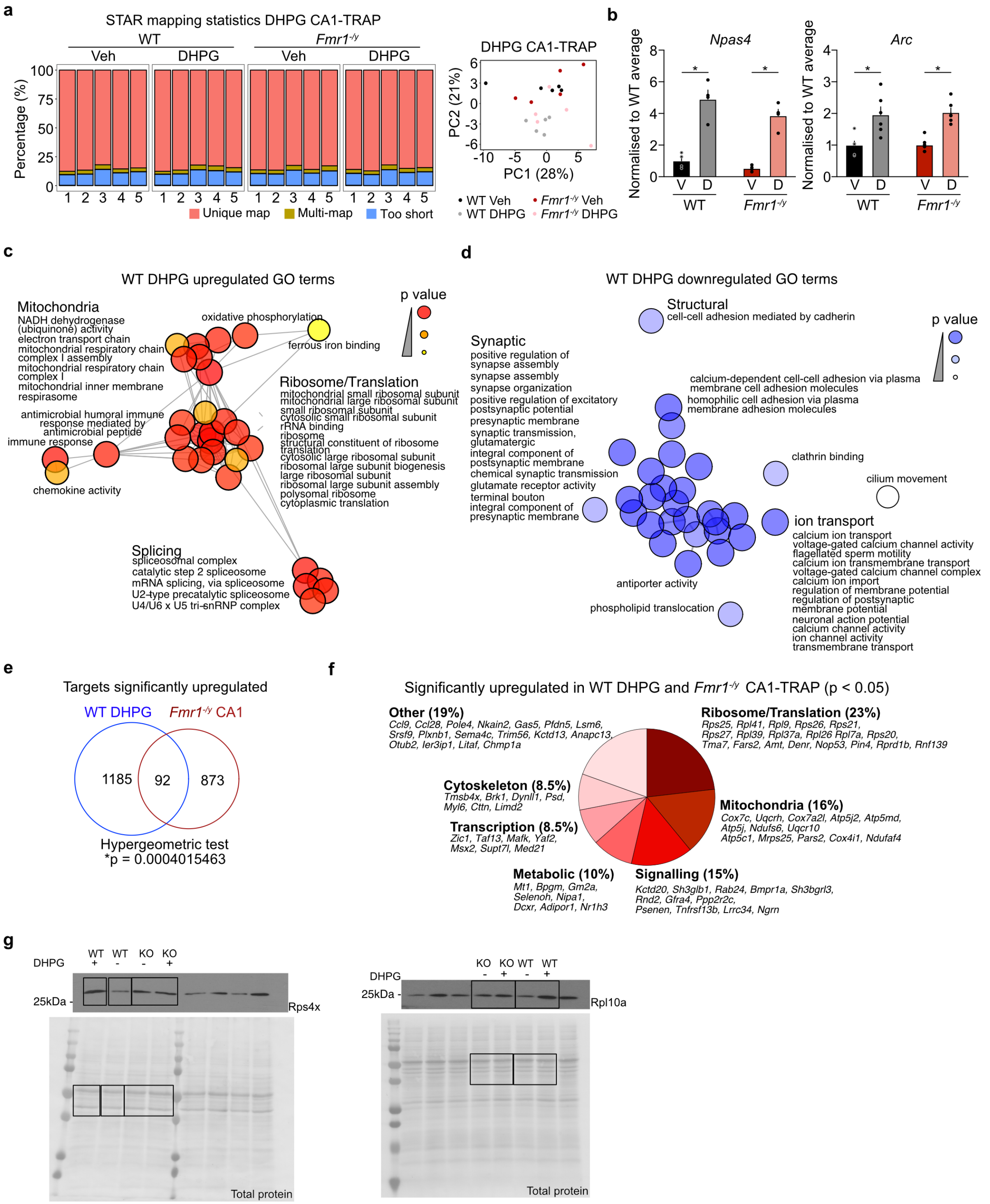


**Supplementary Figure 2. LTD TRAP-seq. (a)** Mapping statistics and PCA plot for Veh and DHPG treated WT and *Fmr1*$^{-/y}$ samples are shown. **(b)** DHPG stimulations were performed on additional WT and *Fmr1*$^{-/y}$ hippocampal slices, and CA1-TRAP performed. qPCR analyses validate the upregulation of immediate early genes *Npas4* (Two-way ANOVA treatment $p < 0.0001$, WT $p < 0.0001$, KO $p < 0.0001$) and *Arc* (Two-way ANOVA $p < 0.0001$, WT $p = 0.0002$, KO $p = 0.0002$) in both genotypes. **(c)** GSEA of transcripts upregulated in WT CA1-TRAP shows an enrichment in ribosomal, mitochondrial and splicing terms (adjusted p value

< 0.01). (**d**) GSEA of transcripts downregulated with DHPG in WT reveals enrichment of terms related to synaptic function, structure, and ion transport (adjusted p value < 0.01). (**e-f**) Analysis of significantly upregulated genes both in WT DHPG and *Fmr1*$^{-/y}$ CA1-TRAP ($p < 0.05$) identifies transcripts involved in ribosome/translation, as well as in mitochondria, cytoskeleton and signaling. (**g**) Raw immunoblots and total protein memcode staining used to quantify Rpl10a and Rps4x expression in **Fig. 2** are shown. As samples are loaded blind to genotype, the arrangement is random and must be re-ordered for the main figure with spaces denoting lanes that are not run next to each other.

**Supplementary Figure 3**

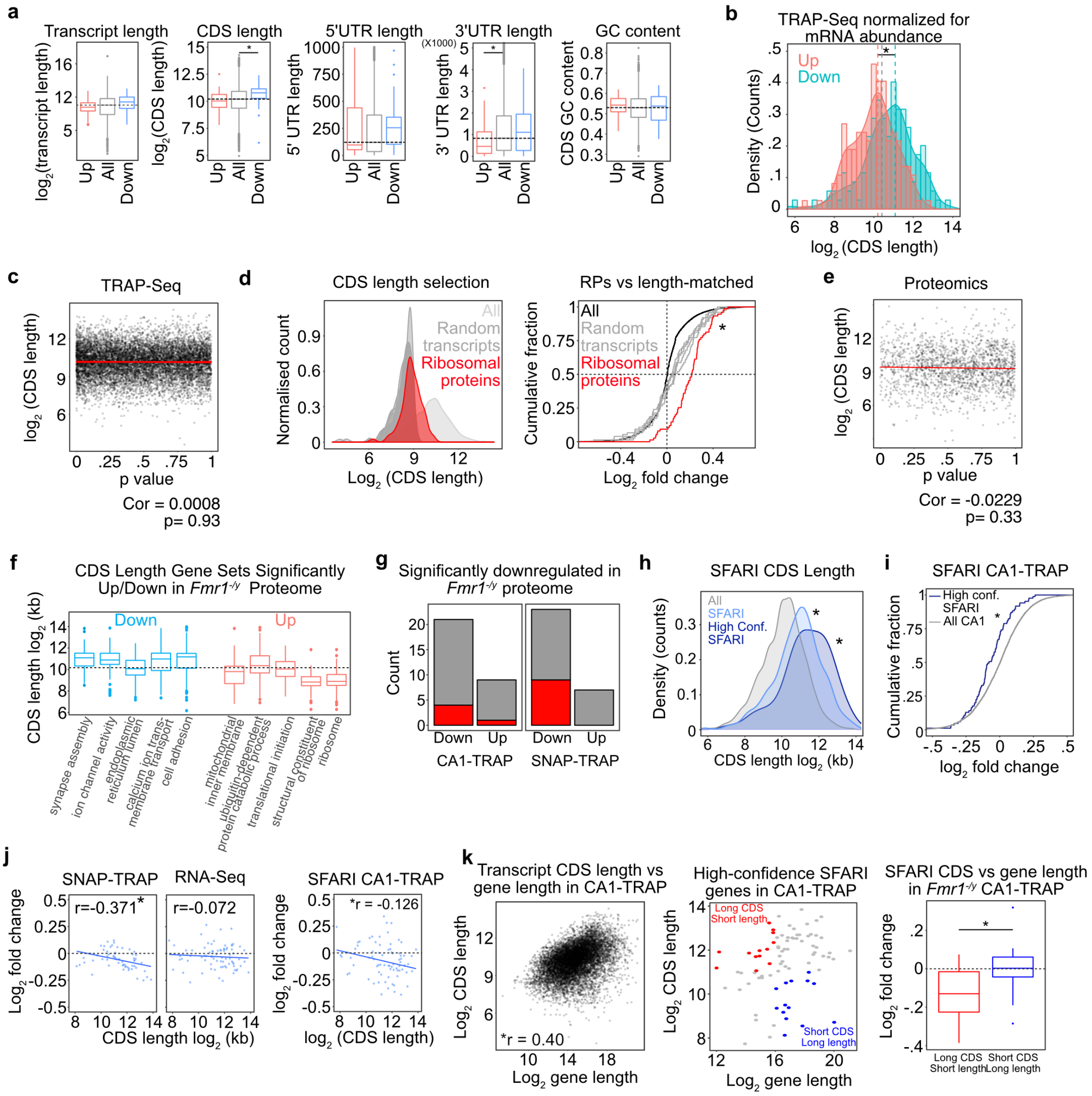


**Supplementary Figure 3. A length-dependent translation imbalance is present in *Fmr1*$^{-/y}$ neurons. (a)** The significantly changed population in CA1-TRAP shows the same length-dependent imbalance in CDS length and 3'UTR length, with a trend towards an imbalance in total transcript length and 5'UTR length. GC content exhibits no difference (CDS down vs average p = 0.012, 3'UTR up versus average p= 0.0078). (**b**) To test whether the length shift in translating mRNAs in *Fmr1*$^{-/y}$ neurons was due to a difference in mRNA abundance, DESeq2 normalized counts for each transcript identified in the TRAP was divided by DESeq2 normalized counts in the total population after CA1 gene filtering. Analysis of translating mRNAs normalized to total shows the same increased length in the downregulated population, indicating this effect is not due to a change in overall transcript abundance (two sample z test, Up vs all z = -1.94, p = 0.051, Down vs all z = 4.17, p = 3.039 X $10^{-5}$). (**c**) An inherent bias in RNA-seq analyses can result in identification of more significant differences in long transcripts [53]. To investigate whether this bias might contribute to the observed length shift in *Fmr1*$^{-/y}$ CA1-TRAP, we performed a correlation analysis between p-value vs CDS length. This analysis reveals no significant correlation, indicating length bias does not explain the changes observed in *Fmr1*$^{-/y}$ CA1-TRAP (Pearson's correlation test r = 0.0008, p = 0.93). **(d)** Comparison of the upregulation of RPs to 5 randomly generated gene sets of the same length shows the elevation in RPs is more significant than what would be predicted from length (RP versus total two sample z test p = 1.03 X $10^{-29}$, short transcripts versus RPs highest adjusted p = 4.35X$10^{-5}$). (**e**) A correlation analysis of the *Fmr1*$^{-/y}$ proteomics dataset reveals no significant correlation between p-value and CDS length (Pearson's correlation test r = -0.0229, p = 0.33). **(f)** Comparison of the average CDS length of mRNAs encoding proteins in the most over- and underexpressed gene sets in the *Fmr1*$^{-/y}$ proteome shows how the CDS length bias in the *Fmr1*$^{-/y}$ translating population can manifest in functionally relevant changes in synaptic protein makeup. **(g)** There is a significant overlap between the targets significantly downregulated in the *Fmr1*$^{-/y}$ proteome and those significantly downregulated in SNAP-TRAP and CA1-TRAP populations (threshold p < 0.05, hypergeometric test p = 7.71 X $10^{-6}$, p = 0.034, SNAP-TRAP and CA1-TRAP respectively). (**h**) Autism risk factors identified as high-confidence by SFARI exhibit significantly longer CDS lengths compared the average TRAP population (two sample z test, all vs SFARI z = 11.234, *p < 2.2X$10^{-16}$, all vs high SFARI z = 6.2066 *p = 5.415X$10^{-10}$). (**i**) Similar to the SNAP-TRAP population, SFARI targets are downregulated in the CA1-TRAP population (two sample z test, z = -4.20, p = 2.619X$10^{-5}$). (**j**) A significant negative correlation seen between CDS length and expression of SFARI transcripts in the *Fmr1*$^{-/y}$ SNAP-TRAP population (Pearson's correlation test r = -0.3714, *p = 0.003). This correlation is not observed in the total *Fmr1*$^{-/y}$ transcriptome (Pearson's correlation test r = -0.17, p = 0.593). Similar to the SNAP-TRAP population, SFARI targets in the *Fmr1*$^{-/y}$ CA1-TRAP population exhibit a significant negative correlation between transcript length and expression (Pearson's correlation test r = -0.126, p = 0.0001). **(k)** Gene length and transcript CDS length are correlated in the CA1 TRAP-seq population (Pearson's correlation test r = 0.40, p < 2.2 X $10^{-16}$). However, a comparison between long genes with short CDS transcripts versus short genes with long CDS transcripts within the SFARI population reveals the differential expression in *Fmr1*$^{-/y}$ CA1-TRAP is driven by the CDS length of the transcript (Wilcox rank sum test p = 0.03766).

**Supplementary Figure 4**

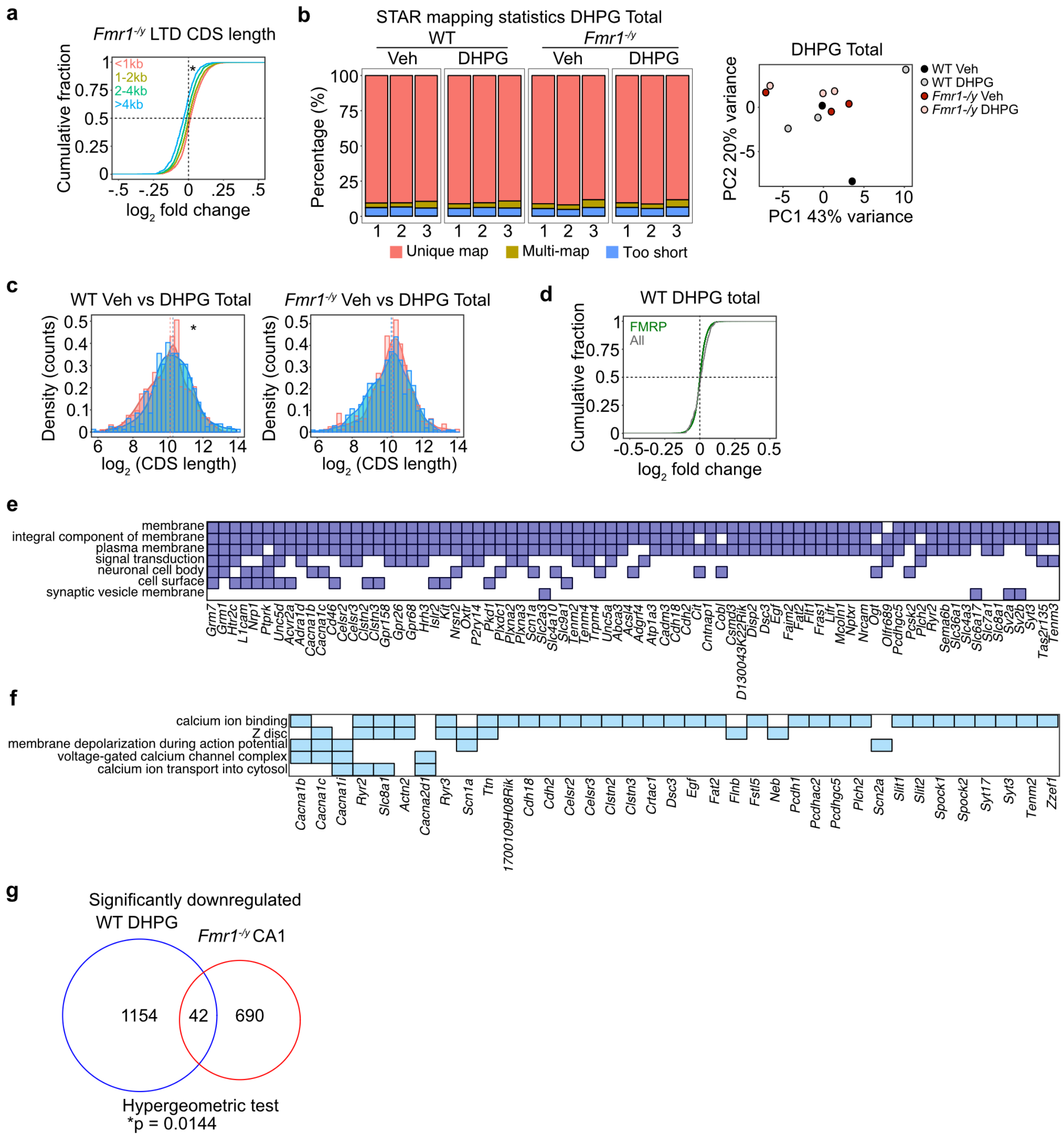


**Supplementary Figure 4. DHPG induces a length-dependent shift in translation in CA1 pyr neurons.** **(a)** Although there are few significantly different changes evoked with DHPG in the *Fmr1*$^{-/y}$ CA1-TRAP, a small but significant length-dependent shift is seen in the population (KS test <1kb vs >4kb p = 3.863 x $10^{-11}$). **(b)** Mapping statistics, PCA plot, and DESeq2 results for the DHPG transcriptome RNA-seq samples are shown (n = 3 littermate pairs). **(c)** Analysis of the total transcriptome of WT and *Fmr1*$^{-/y}$ hippocampus shows that DHPG does not cause the same length-dependent shift as seen in the CA1-TRAP. In fact, there is a slight significant increase in length in the DHPG-upregulated population in WT, which is opposite to the CA1-TRAP fraction (two sample z test, WT: up z = -2.758, *p = 0.0058, down z = 0.2599 p = 0.7949, KO: up z = 0.64304 p = 0.52019, down z = -1.023 p = 0.30638). **(d)** Analysis of FMRP targets in the total transcriptome of DHPG-treated WT slices shown no significant shift in expression (two sample z test, z = -2.039, p = 0.041). **(e)** Clustering of GO terms significantly enriched in the population of transcripts downregulated in WT DHPG is driven by many synaptic elements including large groups of cadherins/protocadherins (*Pcdhac2, Pcdh1, Celsr3, Celsr2, Cdh18, Cdh2, Pcdhgc5*, etc.) and cell adhesion molecules (*L1cam, Nrcam, Focad, Cadm3*, etc.). (**f**) The ion channel cluster downregulated in WT DHPG is driven by multiple targets that are involved in calcium regulation downstream of $mGlu_{1/5}$ activation, including voltage-gated calcium channel transcripts (*Cacna1b, Cacna1c, Cacna1i, Cacna1ad2*) and ryanodine receptors (*Ryr2 and Ryr3*). (**g**) Comparison of the populations significantly downregulated in WT DHPG and in *Fmr1*$^{-/y}$ CA1-TRAP reveals a significant overlap of 42 transcripts (Hypergeometric test, $p = 0.0144$).

**Supplementary Figure 5**

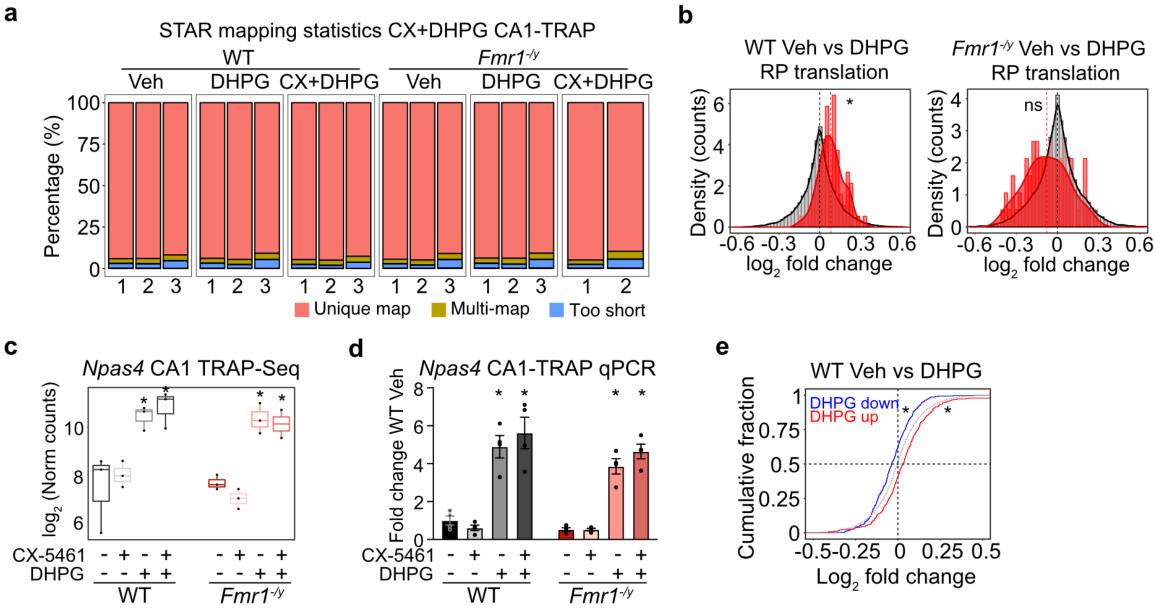


**Supplementary Figure 5. CX-5461 inhibits the length-dependent translation shift in DHPG-treated CA1 pyr neurons. (a)** Mapping statistics for the CX+DHPG TRAP-seq samples are shown. (**b**) Replicating the results of our first DHPG CA1-TRAP experiment, the CX+DHPG dataset shows a significant increase in RP expression in vehicle treated WT after stimulation with DHPG (two sample z test z = 3.0385, p = 0.0023). Also replicating our first experiment, there is no significant increase in RP expression in DHPG-treated *Fmr1*$^{-/y}$ CA1-TRAP (two sample z test z = 1.794207, p = 0.0727). **(c)** TRAP-seq shows that CX-5461 treatment does not alter the upregulation of immediate early gene *Npas4* with DHPG stimulation in either WT or *Fmr1*$^{-/y}$, indicating no change in responsiveness to mGlu$_{1/5}$ activation. (**d**) A qPCR analysis of additional experiments validate these results (Two-way ANOVA treatment p <0.0001, WT Veh vs CX FDR = 0.052, Veh vs DHPG *FDR = 0.0111, Veh vs CXDHPG *FDR 0.0117, *Fmr1*$^{-/y}$ Veh vs CX FDR = 0.3396, Veh vs DHPG *FDR = 0.0019, Veh vs CXDHPG *FDR = 0.0019). **(e)** Transcripts identified as significantly upregulated/downregulated in the first DHPG CA1-TRAP experiment are significantly upregulated/downregulated with DHPG in the WT CA1-TRAP population in the CX+DHPG dataset (two sample z test, LTD up: z = 4.634, *p = 3.588X10$^{-6}$, LTD down: z = -4.7643, *p = 1.895X10$^{-6}$).
